# SHIP2-SRC-β-catenin signaling axis sustains thymidylate synthase expression and promotes fluoropyrimidine resistance

**DOI:** 10.64898/2026.06.05.730406

**Authors:** Abdulrahman El Sayed, Abdelhalim Azzi

**Affiliations:** Laboratory of Lipids and Chronobiology, International Institute of Molecular Mechanisms and Machines (IMol), Polish Academy of Sciences, 00-783, Warsaw, Poland

## Abstract

Fluoropyrimidine-based chemotherapies, including 5-fluorouracil (5-FU) and floxuridine (FuDR), are widely used in cancer treatment, but their efficacy is limited by adaptive resistance driven by TYMS upregulation. The upstream mechanisms controlling TYMS expression remain poorly defined. Here, we identify INPPL1 (SHIP2) as a critical regulator of TYMS expression and fluoropyrimidine response in breast cancer cells. We show that SHIP2 enhances basal and drug-induced TYMS expression at the transcriptional level independently of its phosphatase activity. Mechanistically, SHIP2 increases SRC levels and nuclear accumulation of β-catenin, driving TYMS expression. Inhibition of SRC or β-catenin suppresses TYMS induction and restores sensitivity to FuDR. Importantly, SHIP2 rewires TYMS regulation from a P53-dependent program to a β-catenin-driven pathway, enabling sustained TYMS expression under chemotherapeutic stress. Consistent with this model, differential sensitivity to SHIP2 depletion correlates with baseline TYMS levels across cell lines. Analysis of patient cancer datasets reveals that high INPPL1 expression correlates with increased TYMS levels and poor clinical outcomes. These findings identify SHIP2 as a non-canonical regulator of TYMS and a potential therapeutic target to overcome fluoropyrimidine resistance.

## INTRODUCTION

Fluoropyrimidine-based chemotherapies, including 5-fluorouracil (5-FU) and its metabolite floxuridine (FuDR), remain widely used for the treatment of colorectal, gastrointestinal, and breast cancers. Floxuridine primarily binds to and inhibits thymidylate synthase (TYMS), thereby disrupting deoxythymidine monophosphate (dTMP) synthesis, inducing nucleotide imbalance, replication stress, and DNA damage [1]. However, acquired resistance frequently limits their clinical efficacy. One of the primary mechanisms of resistance to fluoropyrimidine-based chemotherapies involves adaptive upregulation of TYMS, which restores nucleotide pools and attenuates drug-induced cytotoxicity [2–4]. Nevertheless, the upstream signaling pathways controlling this adaptive response remain poorly defined. The SH2 domain-containing inositol 5-phosphatase 2 (SHIP2) is a lipid phosphatase that hydrolyzes PI (3,4,5) P3 to PI (3,4) P2, which negatively regulates the PI3K/AKT signaling axis [5]. Beyond its catalytic function, SHIP2 also acts as an adaptor protein. Indeed, its interaction with the SRC family kinases promotes sustained ERK activity upon FGF stimulation [6]. Increasing evidence implicates SHIP2 in tumor progression. In fact, elevated SHIP2 expression has been associated with aggressive phenotypes and poor outcomes in multiple malignancies, including esophageal, breast, and colorectal cancers [7–8]. Despite these observations, the role of SHIP2 in chemotherapeutic adaptation, particularly in the regulation of nucleotide metabolism, remains unexplored.

β-catenin is a central effector of canonical Wnt signaling [9]. In concert with other transcription factors, β-catenin acts as a transcriptional co-activator, driving expression of genes involved in proliferation, stemness, and survival [10]. Aberrant stabilization and nuclear accumulation of β-catenin have been implicated in resistance to multiple anticancer therapies, including fluoropyrimidines [11–12]. Given its role as a transcriptional regulator of genes involved in cell cycle progression and metabolic adaptation, β-catenin represents a plausible intermediary protein linking upstream signaling pathways to TYMS expression and drug resistance. In addition, SRC family kinases, such as c-SRC, have been reported to regulate TYMS expression, although the underlying mechanisms remain incompletely understood [4].

Based on these observations, we hypothesized that SHIP2 functions as an upstream regulator of adaptive fluoropyrimidine resistance by coordinating SRC-β-catenin signaling and TYMS expression. Using breast cancer as a model and combining correlative analyses, gain and loss-of-function approaches, and pharmacological perturbation, we identify a previously unrecognized SHIP2-SRC-β-catenin-TYMS signaling axis that promotes resistance to Floxuridine.

We demonstrate that SHIP2 enhances both basal and drug-induced TYMS expression through a phosphatase-independent mechanism and promotes β-catenin stabilization and transcriptional activity. Furthermore, β-catenin activity is required for SHIP2-mediated adaptive responses to fluoropyrimidine treatment, as its inhibition restores cellular sensitivity to floxuridine. Notably, we show that SHIP2 rewires TYMS regulation from a p53-dependent to a β-catenin-dependent pathway and identify SRC kinase as a critical mediator of this pathway. Collectively, our findings establish SHIP2 as a non-canonical regulator of nucleotide metabolism and uncover a signaling mechanism that promotes chemotherapeutic resistance, revealing potential therapeutic vulnerabilities in fluoropyrimidine-treated cancers.

## RESULTS

### SHIP2 promotes TYMS transcription independently of its phosphatase activity

The phosphoinositide 5-phosphatase SHIP2 negatively regulates the PI3K/AKT signaling pathway, it has also emerged as a multifunctional regulator of oncogenic signaling [13]. SHIP2 expression is elevated in multiple tumor types compared to adjacent normal tissues [14]. Despite its negative impact on PI3K/AKT signaling, recurrent SHIP2 upregulation in cancer suggests that it may confer a fitness advantage or facilitate adaptation to adverse environmental conditions, such as metabolic stress or chemotherapy.

To investigate whether SHIP2 is linked to fluoropyrimidine response, we first examined the relationship between INPPL1 (encoding SHIP2) and TYMS expression. Analysis of RNA-sequencing data from the GEPIA2 database [15] revealed a significant positive correlation between INPPL1 and TYMS expression across all cancer types (**Figure 1A**). Moreover, this correlation was preserved when the analysis was restricted to breast cancer samples (**Figure 1B**), suggesting that SHIP2 expression is associated with elevated TYMS levels in both pan-cancer and breast-specific contexts. To determine whether SHIP2 directly regulates TYMS expression, we used lentiviral-mediated overexpression of wild-type (WT) SHIP2 or a phosphatase-dead (PD) SHIP2 mutant in MDA-MB-231 breast cancer cells. As shown in Figure 1C, immunoblot analysis revealed that both WT and PD SHIP2 overexpression resulted in a marked increase in TYMS protein levels compared to control cells. Consistent with the known phosphoinositide 5-phosphatase activity of SHIP2, phosphorylation of AKT was significantly reduced in WT SHIP2-expressing cells but not in PD SHIP2 cells, suggesting that the increase in TYMS levels is independent of SHIP2 lipid phosphatase activity (**Figure 1 C-E**). TYMS protein levels are regulated at both transcriptional and translational levels [4,16]. Thus, we examined whether the increase in TYMS observed upon overexpression of WT and PD SHIP2 is due to increased transcription. Indeed, quantitative RT-PCR analysis demonstrated that TYMS mRNA levels were significantly elevated in both WT and PD SHIP2-expressing cells (**Figure 1 F-G**), indicating that SHIP2 promotes TYMS expression at the transcriptional level independently of its lipid phosphatase activity.

**Figure 1:**
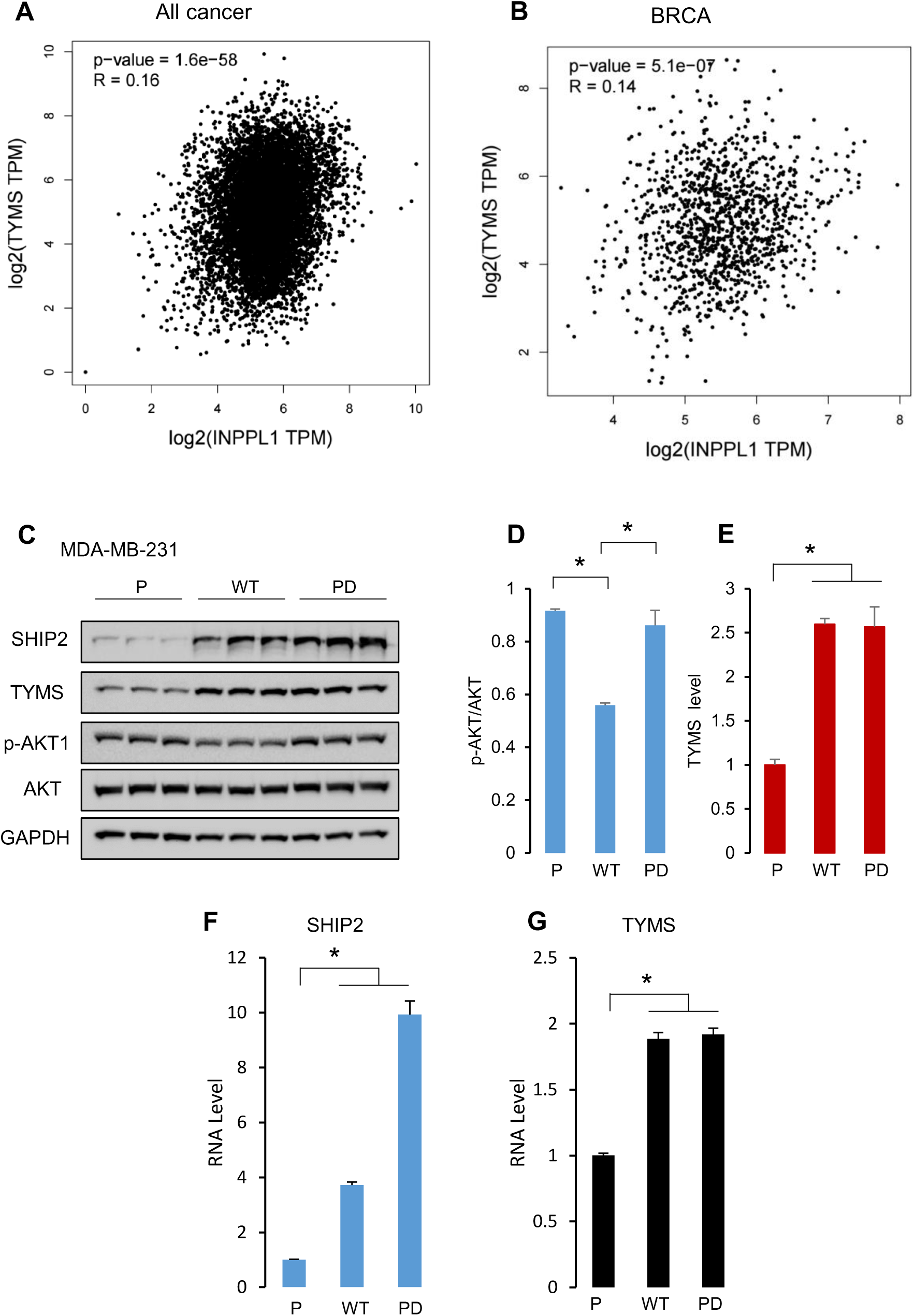
SHIP2 positively correlates with TYMS and promotes its transcription. **(A-B)** SHIP2 transcript levels positively correlate with TYMS RNA levels in cancer. Scatterplot displaying results of Spearman correlation between SHIP2 and TYMS expression in all cancers when compared to non-tumorigenic counterparts (A) and in breast cancer (B). **(C)** Lysates from MDA-MB-231 parental cells or stably overexpressing WT or PD SHIP2 were stained with the indicated antibodies. n=3 biological replicates per genotype. **(D-E)** Bar graphs showing the results of densitometric analysis of AKT1 phosphorylation and TYMS protein levels under the conditions obtained in C. The data are presented as the means ±SEMs. ANOVA, multiple comparisons: Dunnett test. p-AKT1: F (2, 6) = 33,23 P <0.005. TYMS: (F (2, 6) = 52.58, P <0.0005). **(F-G)** Bar graph showing Tyms mRNA levels in MDA-MB-231 parental or stably overexpressing WT or PD SHIP2. The data are presented as the means ±SEMs. ANOVA, multiple comparisons: Dunnett test. TYMS (F (2, 6) = 164, P <0.0001).

### SHIP2 enhances TYMS induction in response to 5-FU and FuDR

TYMS expression and protein levels are known to increase in response to fluoropyrimidine treatment as part of an adaptive resistance mechanism [2, 4]. We therefore asked how the increase in WT SHIP2 and PD SHIP2 levels in MDA-MB-231 cells affects TYMS protein levels after 24 hours of 5-FU treatment. As indicated in **Figure 2A**, TYMS protein levels increased in all conditions; however, the magnitude of induction was markedly higher in WT and PD SHIP2-overexpressing cells (**Figure 2 A-B**).

**Figure 2:**
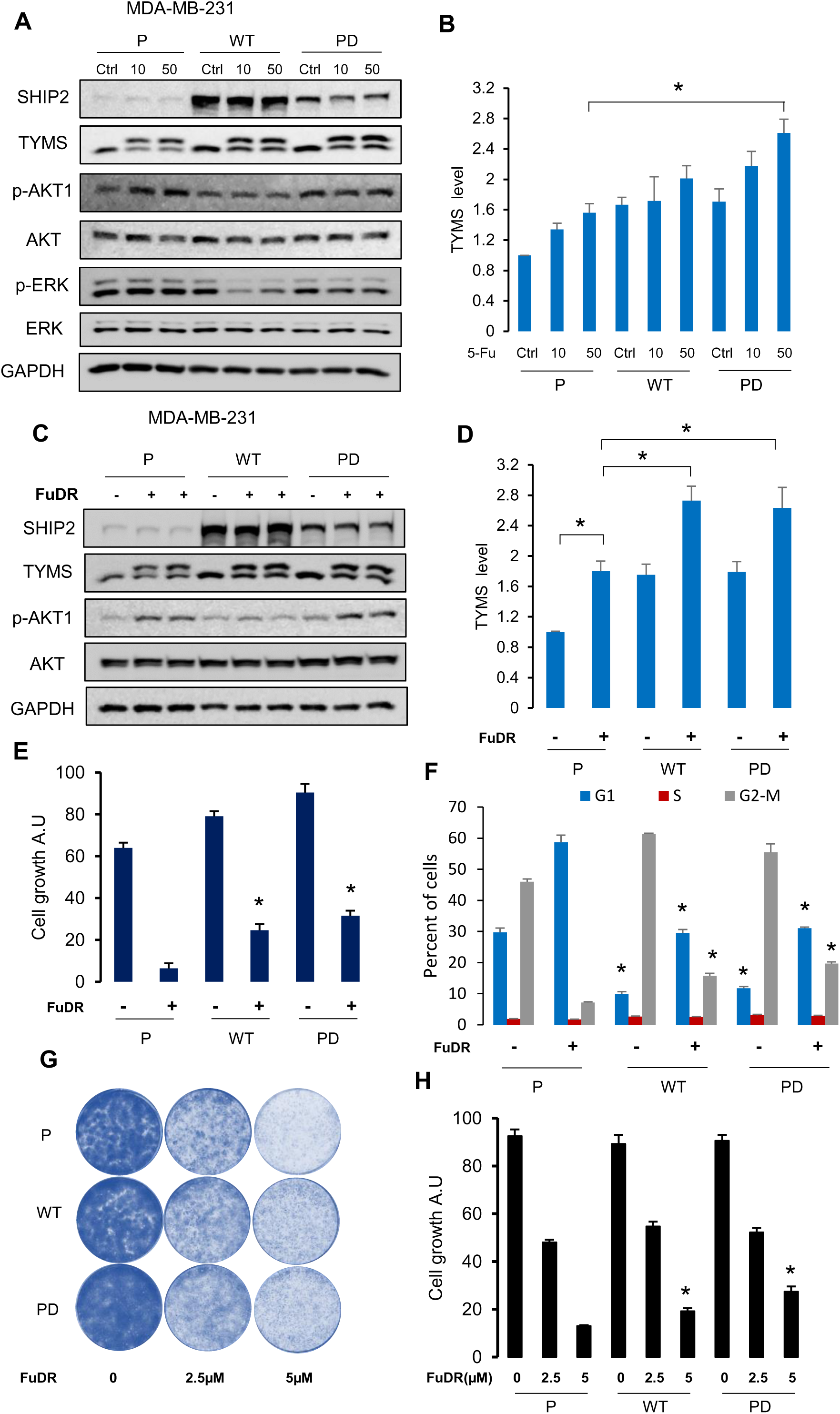
Elevated SHIP2 enhances the 5-FU-mediated increase in TYMS protein and protects against 5-FU-mediated cell death. (A, C) Lysates from MDA-MB-231 parental cells or stably overexpressing WT or PD SHIP2 control or treated with 5-FU or FuDR for 24 hours, then stained with the indicated antibodies. n=3 biological replicates per genotype. (B, D) Bar graphs showing the results of densitometric analysis of TYMS protein levels under the conditions obtained in A and C. The data are presented as the means ±SEMs. ANOVA, multiple comparisons: Dunnett test. (B) TYMS: F (8, 18) = 7.68, P <0.001. (D) TYMS: F (5, 24) = 11.4, P <0.001. (**E**) Bar graph showing cellular ATP content in MDA-MB-231 parental cells or stably overexpressing WT or PD SHIP2 control or treated with FuDR for 72 hours. Results expressed as a percentage of control. Data are the means ± SEM from 3 independent experiments. One-way ANOVA, F(5, 24) = 132.5. (**F**) Bar graphs showing the percentage of MDA-MB-231 parental, WT, and PD SHIP2 cells at different phases of the cell cycle after 72 hours of treatment with 50µM FuDR. Data are presented as the means ±SEMs. ANOVA G1 (F (5, 12) = 193.1, **P < 0.001). S (F (5, 12) = 9.01, **P<0.005). G2-M F (5, 12) = 334, **P<0.0001). (**G-H**) Clongenic assay of MDA-MB-231 parental, WT, and PD SHIP2 cells treated with the indicated concentrations of FuDR. Representative images from three biological replicates are shown for each condition. ANOVA, multiple comparisons: Dunnett test. MDA-MB-231: F (11, 60) = 368, P<0.0001.

The metabolism of 5-FU leads to the formation of 5-fluorouridine and 5-fluorodeoxyuridine (Floxuridine), which are then incorporated into RNA and DNA, respectively, causing damage to these nucleic acids [1, 17]. Since 5-fluorodeoxyuridine is the main interacting molecule and inhibitor of TYMS protein, we examined in more detail the impact of FuDR, allowing us to isolate the effects of direct thymidylate synthase inhibition from RNA-directed toxicity. MDA-MB-231 control, WT, and PD cells were treated with 5-fluoro-2′-deoxyuridine (FuDR) for 24 hours. Consistent with the results obtained with 5-FU, FuDR treatment induced TYMS protein accumulation in all cells; however, WT and PD SHIP2-expressing cells exhibited significantly higher TYMS levels than the controls (**Figure 2 C-D**). We next assessed whether SHIP2-mediated increase of TYMS confers functional resistance to thymidylate synthase inhibition. In the short-term growth assay using cellular ATP content, parental cells displayed markedly lower ATP levels under basal conditions and after 72 hours of treatment with FuDR (**Figure 2 E**). Moreover, cell cycle analysis after 72 hours of FuDR exposure revealed that SHIP2 overexpression reduced FuDR-induced G1 arrest. Indeed, both WT and PD SHIP2-expressing cells exhibited a lower proportion of G1-phase cells than parental cells (**Figure 2F**), indicating that SHIP2 supports continued cell cycle progression under thymidylate synthase inhibition. Furthermore, in long-term colony formation assays, WT and PD SHIP2-expressing MDA-MB-231 cells exhibited increased colony numbers following high FuDR concentration compared to parental cells (**Figures 2G-H**). Together, these data demonstrate that increased SHIP2 protein levels confer resistance to Floxuridine. However, long-term FuDR treatment results in greater resistance in cells that overexpress the phosphatase dead SHIP2 mutant. Although cells overexpressing the WT form of SHIP2 showed a similar trend, the magnitude of resistance was lower, suggesting that downregulation of PI3K/AKT by the phosphatase-active form of SHIP2 may partially limit resistance during prolonged FuDR treatment.

### SHIP2 depletion alters TYMS expression and sensitizes cells to FuDR in a context-dependent manner

Our data above indicate that high levels of SHIP2 protein confer resistance to FuDR-mediated growth arrest, independent of its lipid phosphatase activity. To determine whether SHIP2 is required for TYMS regulation and cellular response to Floxuridine, SHIP2 was stably depleted using shRNA in both MCF7 and MDA-MB-231 cells. As shown in **Supplementary Figure 1,** this approach leads to a marked downregulation of SHIP2 protein in both cell lines. Interestingly, SHIP2 depletion resulted in a pronounced reduction in TYMS protein levels under basal conditions, with a more pronounced effect in MCF7 cells. To examine whether this regulatory relationship is preserved in other cell models, we assessed the impact of its knockdown in HEK293T cells. Consistent with our observations in breast cancer cells, SHIP2 depletion in HEK293T cells resulted in a significant decrease in TYMS protein levels (**Supplementary Figure 1 C**). These findings suggest that SHIP2 is required to maintain high levels of TYMS protein across distinct cellular backgrounds.

We next examined how SHIP2 depletion affects cellular response to FuDR treatment. As shown in Figure 3 A-B, the decrease of TYMS protein persists upon FuDR treatment. Indeed, Sh control cells treated with FuDR exhibited the expected marked increase in TYMS protein, whereas SHIP2-depleted cells failed to display an increase in TYMS protein (**Figure 3 A-B, 3D, 3F**). Moreover, this experiment further confirmed that SHIP2-mediated regulation of TYMS levels does not depend on the PI3K/AKT signaling, as SHIP2 depletion leads to a markedly higher AKT phosphorylation, especially in MCF7 cells. Furthermore, these data were also validated using a separate ShRNA targeting SHIP2 in MDA-MB-231 cells (**Supplementary Figure 2 A-B**).

**Figure 3:**
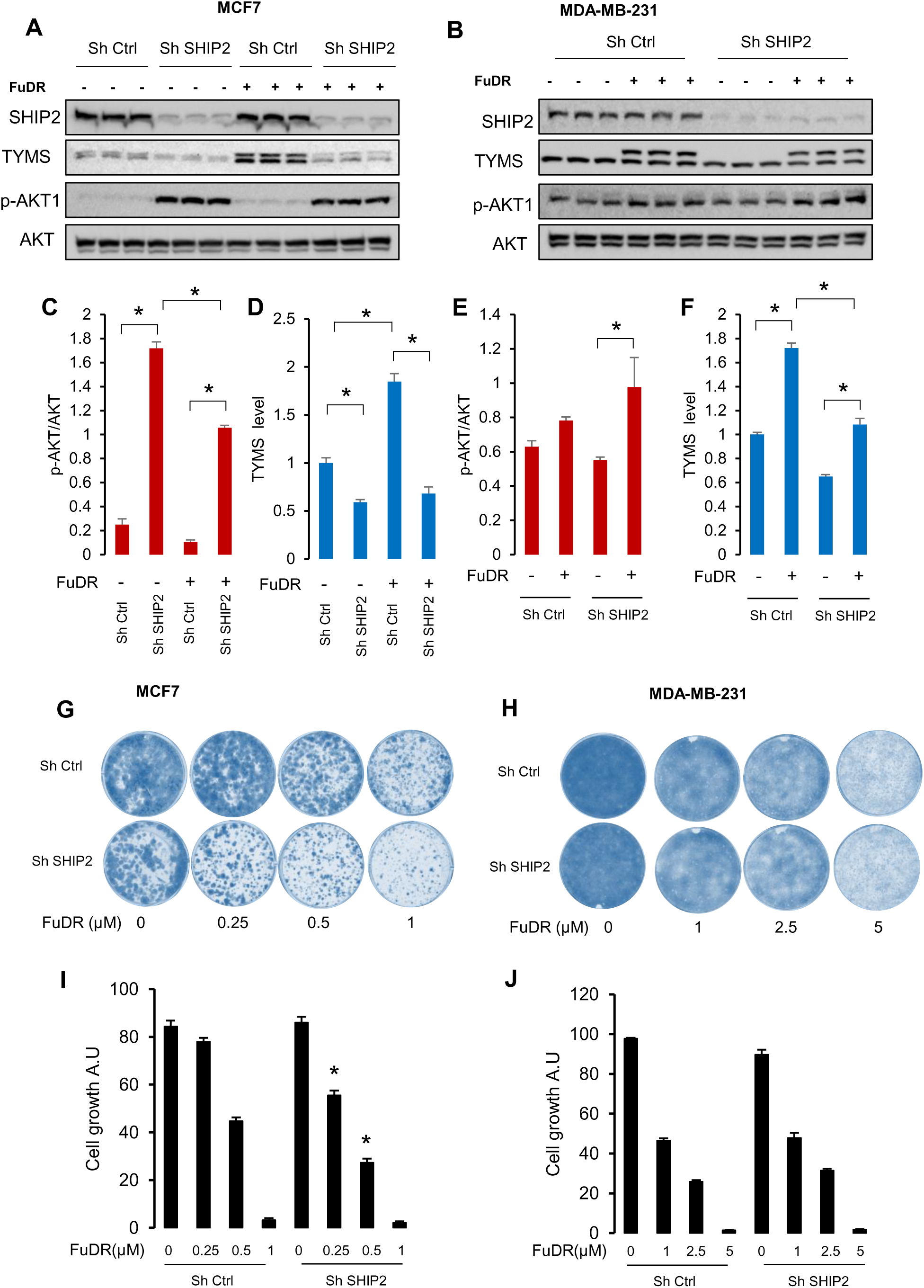
SHIP2 depletion alters cellular response to FuDR. (**A-B**) Lysates of MDA-MB-231 and MCF7 cells stably expressing Sh control or small hairpin against SHIP2, treated or not with 50µM 5-FuDR for 24 hours, were stained with the indicated antibodies. (**C-F**) Bar graphs showing the results of densitometric analysis of AKT1 phosphorylation and TYMS protein levels under the conditions obtained in A and B. The data are presented as the means ±SEMs. ANOVA, multiple comparisons: Dunnett test. MCF7: p-AKT1 F (3, 8) = 390.6, P <0.0001, TYMS F (3, 8) = 83.3, P <0.0001. MDA-MB-231: p-AKT1 F (3, 8) = 4.43, P<0.05. TYMS F (3, 8) = 154.1, P<0.001. (**E-H**) Clongenic assay of MDA-MB-231 and MCF7 cells stably expressing Sh control or small hairpin against SHIP2 treated with the indicated concentrations of FuDR. Representative images from three biological replicates are shown for each condition. ANOVA, multiple comparisons test. MCF7: F (7, 88) = 377.5, P<0.0001. MDA-MB-231: F (7, 88) = 682.8, P<0.0001.

Together, these findings indicate that SHIP2 is required for the FuDR-induced increase in TYMS protein levels and suggest that loss of SHIP2 compromises a key adaptive resistance mechanism to Floxuridine treatment.

We next examined how SHIP2 depletion functionally affects the response to FuDR. Using a colony-forming assay, we monitored the impact of SHIP2 depletion on long-term cell survival. These experiments revealed that the impact of SHIP2 depletion is cell-type-dependent. Indeed, while we observed a significant decrease in colony-forming ability in MCF7 cells lacking SHIP2 relative to ShRNA control cells, no marked difference was observed in MDA-MB-231 cells after treatment with increasing concentrations of FuDR (**Figure 3 G-J**). To explain this discrepancy, we compared basal TYMS expression levels between the two cell lines. As indicated in Supplementary Figure 3, MDA-MB-231 cells displayed a markedly higher basal TYMS protein levels relative to MCF7 cells. Although SHIP2 knockdown reduced TYMS levels in MDA-MB-231 cells, residual TYMS expression remained substantially higher than that observed in SHIP2-depleted MCF7 cells (**Figure 3 A-B**). These findings suggest that the differential impact of SHIP2 depletion on drug sensitivity may reflect intrinsic differences in baseline TYMS abundance, with MDA-MB-231 cells maintaining TYMS levels above the threshold required to sustain nucleotide synthesis under fluoropyrimidine stress.

Importantly, overexpression of either wild-type or phosphatase-dead SHIP2 in MDA-MB-231 cells further increased TYMS levels and conferred enhanced resistance to FuDR, indicating that SHIP2 functions as an amplifier of TYMS expression in a context-dependent manner (**Figure 2**).

Another possibility that might explain the differential sensitivity of these two cell lines to FuDR following SHIP2 depletion is their p53 status. Indeed, MCF7 cells express the wild-type form of p53, whereas MDA-MB-231 cells express a mutant gain-of-function form of p53 [18]. We therefore examined whether p53-dependent stress responses contribute to this differential sensitivity. We specifically analyzed the expression of four well-characterized p53-responsive genes involved in cell cycle control, DNA damage response, oxidative stress regulation, and cell survival: CDKN1A (p21), GADD45, SESN2, and BIRC5 [19]. As shown in **Supplementary Figure 4,** FuDR treatment modulated the expression of these genes in a gene- and cell-type-specific manner upon SHIP2 depletion. Indeed, the transcript levels of CDKN1A and GADD45 were comparable between control and SHIP2-depleted cells, suggesting that SHIP2 loss does not broadly impair p53-mediated cell cycle arrest or DNA damage signaling. In contrast, SESN2 expression was markedly elevated in SHIP2-depleted cells, suggesting an enhanced oxidative stress and increased reactive oxygen species accumulation following SHIP2 loss. Interestingly, BIRC5 expression was significantly reduced in SHIP2-depleted MCF7 cells but increased in SHIP2-depleted MDA-MB-231 cells, suggesting divergent survival responses that may reflect differences in p53 functionality.

Taken together, these data suggest that SHIP2 depletion sensitizes cells to Floxuridine in a context-dependent manner, potentially through combined suppression of TYMS expression and differential activation of p53 and stress-responsive survival pathways.

### SHIP2 promotes nuclear accumulation of active β-catenin

β-Catenin is a central effector of the canonical Wnt signaling pathway and plays a key role in regulating cell proliferation, survival, and stemness. Aberrant β-catenin activation has been reported in several cancers and is also associated with resistance to chemotherapy and radiotherapy [20]. In the context of fluoropyrimidine treatment, β-catenin regulation appears to be highly context-dependent. Indeed, in mice, 5-FU induces hair loss and blocks β-catenin nuclear translocation [21]. In colorectal cancer cells and squamous cell carcinoma, it was reported that resistance to 5-FU is associated with increased activity of the Wnt/β-catenin signaling pathway [12, 22]. Interestingly, increased Wnt/β-catenin signaling enhances the enrichment of cancer stem cells (CSCs) [11]. Based on these observations, we examined whether the modulation of SHIP2 levels affects β-catenin status. In MDA-MB-231 cells, FuDR treatment moderately reduces active and total β-catenin protein levels in control, WT SHIP2, and PD SHIP2 cells (**Figure 4 A-B**). However, levels of active (non-phosphorylated) and total β-catenin were significantly higher in WT and PD SHIP2-overexpressing cells under basal conditions and remained elevated following FuDR treatment compared with control cells (**Figure 4 A-B**). Conversely, SHIP2 depletion in both MCF7 and MDA-MB-231 cells resulted in a moderate decrease in active β-catenin levels under basal conditions, which became more pronounced and significant following FuDR exposure (**Figure 4 C-F**). We next examined whether SHIP2 depletion affects the subcellular distribution of β-catenin. As shown in Figure 4 G-H, cellular fractionation analyses revealed that SHIP2 depletion reduced nuclear levels of active β-catenin in both cell lines. In contrast, MDA-MB-231 cells stably expressing WT or PD SHIP2 showed increased nuclear accumulation of active β-catenin (**Figure 5 A-B**), a phenomenon commonly associated with transcriptional regulation in Wnt-responsive contexts [20].

**Figure 4:**
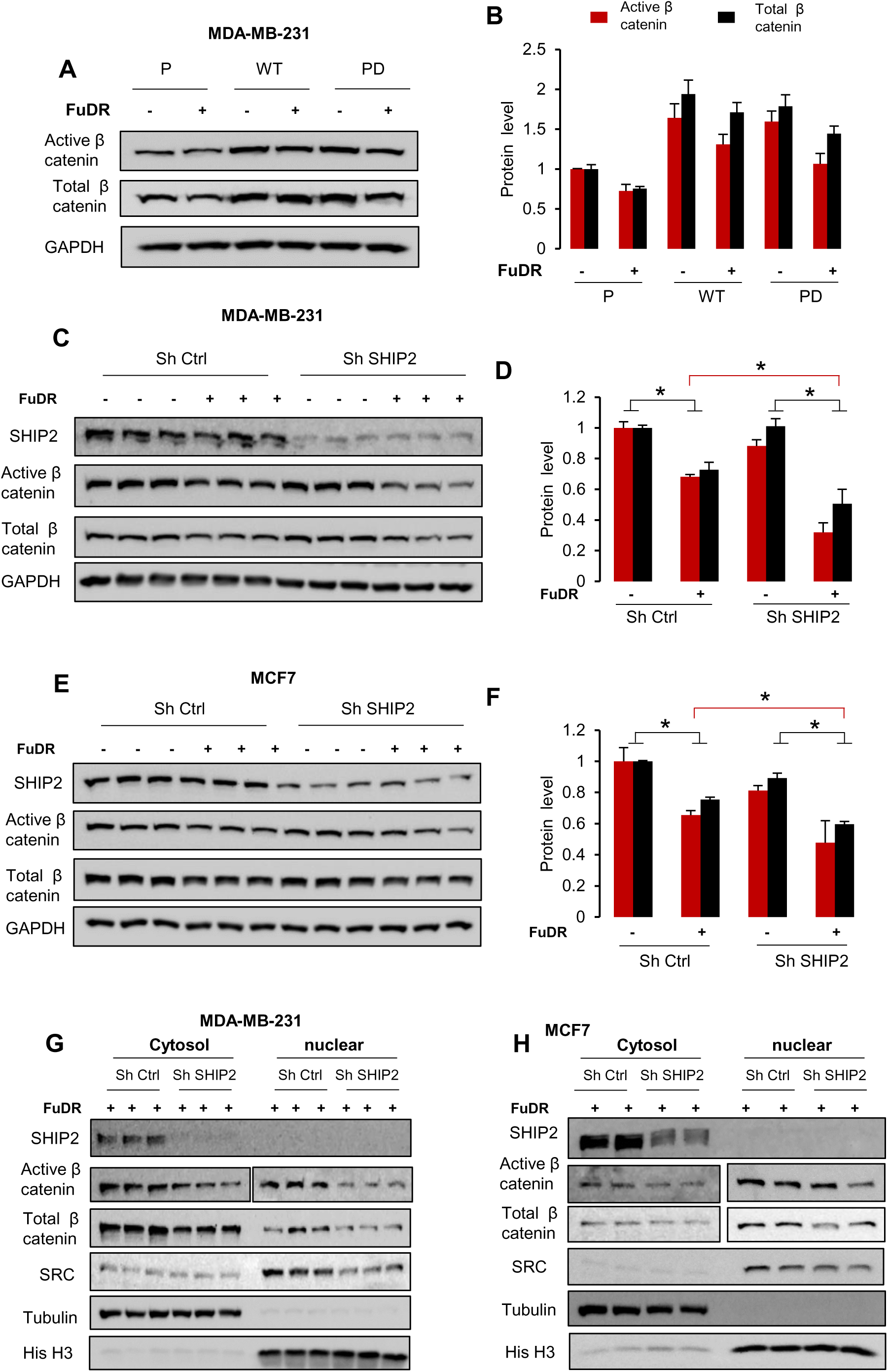
SHIP2 modulates β-catenin levels in response to FuDR-mediated cellular stress. (**A, C, E**) Lysates of MDA-MB-231 and MCF7 cells stably expressing Sh control or small hairpin against SHIP2, and MDA-MB-231 overexpressing WT or PD SHIP2 treated or not with 50µM 5-FuDR for 24 hours, were stained with the indicated antibodies. (**B, D, F**) Bar graphs showing the results of densitometric analysis of active and total β-catenin levels under the conditions obtained in A, C, and E. The data are presented as the means ±SEMs. ANOVA, multiple comparisons: Tukey’s multiple comparisons test. **B**: active β-catenin F (5, 12) = 8.7, P <0.005. Total β-catenin F (5, 12) = 16.85, P<0.0001. (**D)** Active β catenin F (3, 8) = 49.8, P <0.0001, total β catenin F (3, 8) = 16.6, P <0.001, **(F)** active β catenin (3, 8) = 6.7, P<0.05. Total β-catenin F (3, 8) = 76.1, P <0.05. **(G-H)** Western blot analysis of nuclear and cytosolic levels of active and total β-catenin in MDA-MB-231 cells and MCF7 Sh control and Sh SHIP2 treated with FuDR.

**Figure 5:**
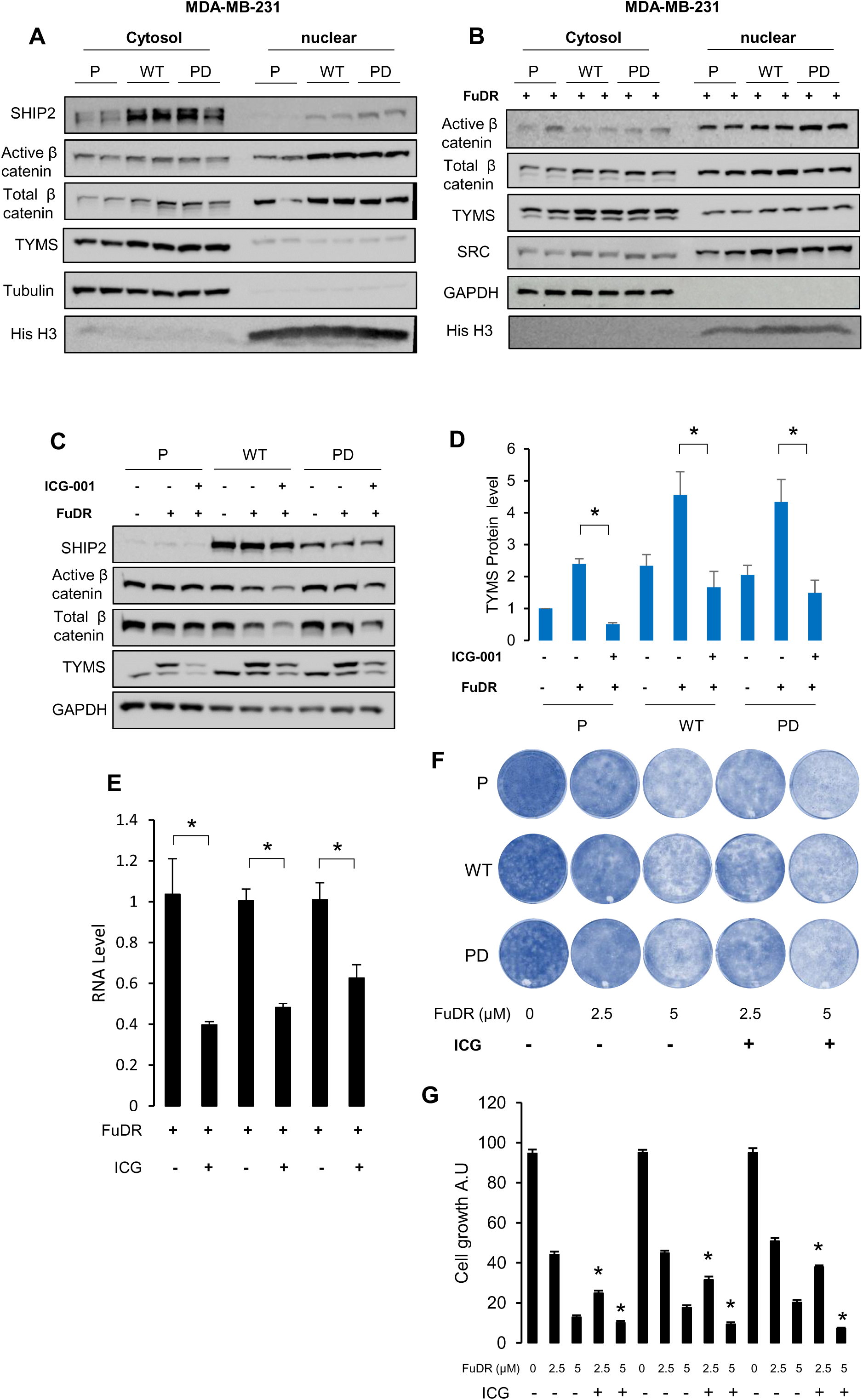
Inhibition of β-catenin reduces TYMS expression and restores sensitivity to FuDR in SHIP2-overexpressing cells. **(A-B)** Western blot analysis of nuclear and cytosolic levels of active and total β-catenin in MDA-MB-231 parental cells and cells overexpressing WT or PD SHIP2 treated with or without FuDR. (a) Control. (b) FuDR. Representative of 3 biological replicates per condition. **(C)** Lysates from MDA-MB-231 parental cells or stably overexpressing WT or PD SHIP2 control or treated with 50 µM FuDR for 24 hours, supplemented or not with 10 µM β-catenin inhibitor (ICG-001), then stained with the indicated antibodies. n=3 biological replicates per condition. **(D)** Bar graphs showing the results of densitometric analysis of TYMS levels under the conditions obtained in C. The data are presented as the means ±SEMs. ANOVA, multiple comparisons: Dunnett test. Parental cells: F (2, 6) = 102.6, P<0.001. WT cells: F (2, 6) = 7.6, P<0.05. PD cells: F (2, 6) = 9.2, P<0.05. (**E**) Bar graph showing TYMS mRNA levels in MDA-MB-231 parental, WT, and PD SHIP2 cells treated with FuDR supplemented or not with 10µM β-catenin inhibitor. Unpaired T-test. *P<0.05. (**F-G**) Clongenic assay of MDA-MB-231 parental, WT, and PD SHIP2 cells treated with the indicated concentrations of FuDR supplemented or not with 1µM β-catenin inhibitor. Representative images from 6 biological replicates are shown for each condition. ANOVA, multiple comparisons: Dunnett test. F (20, 105) = 457.05, 1.34E-92.

Together, these data indicate that SHIP2 expression positively correlates with the accumulation of non-phosphorylated (active) β-catenin under both basal conditions and following FuDR treatment. SHIP2 overexpression is associated with sustained levels and nuclear enrichment of active β-catenin, whereas SHIP2 depletion leads to reduced active β-catenin levels, particularly upon FuDR treatment. These findings suggest that SHIP2 modulates β-catenin stability and subcellular distribution, which in turn might contribute to cellular adaptation to fluoropyrimidine treatment.

### β-Catenin inhibition prevents SHIP2-mediated upregulation of TYMS and FuDR resistance

To determine whether β-catenin mediates SHIP2-dependent TYMS regulation, MDA-MB-231 cells overexpressing WT or phosphatase-dead SHIP2 were treated with the β-catenin/CBP interaction inhibitor ICG-001 in combination with FuDR. ICG-001 selectively disrupts the interaction between β-catenin and CREB-binding protein (CBP) without affecting β-catenin/p300 interaction, thereby inhibiting CBP-dependent β-catenin transcriptional activity [23–24]. Inhibition of β-catenin/CBP signaling markedly attenuated the FuDR-induced increase in TYMS protein levels in both WT and PD SHIP2-overexpressing cells (**Figure 5 C-D**). Moreover, combined treatment with FuDR and ICG-001 significantly reduced TYMS mRNA levels, indicating that the increase in TYMS mRNA observed in cells overexpressing WT or PD-SHIP2 is mediated by β-catenin (**Figure 5 E**). Together, these results demonstrate not only that β-catenin activity is required for TYMS transcription in parental cells, but also that SHIP2-mediated increase in TYMS expression depends on β-catenin.

To determine whether this effect specifically involves CBP-dependent transcription rather than classical β-catenin/TCF4 signaling, cells were treated with PNU-74654, an inhibitor that disrupts the β-catenin/TCF4 interaction [25]. In contrast to ICG-001, PNU-74654 had no significant effect on FuDR-induced TYMS protein levels (**Supp Figure 5**). These findings suggest that SHIP2-mediated TYMS upregulation depends on CBP-associated β-catenin transcriptional activity rather than on β-catenin/TCF4-dependent activation of canonical Wnt target genes.

We next examined how β-catenin functionally affects the response to FuDR. As indicated in Figure 5 F-G, combined treatment of β-catenin inhibition and FuDR reversed the resistance conferred by SHIP2 overexpression. Indeed, cells overexpressing WT or PD SHIP2, which normally exhibit greater colony-forming ability under FuDR treatment, displayed reduced survival and growth when β-catenin was concurrently inhibited. These findings provide functional evidence that β-catenin is a critical mediator of SHIP2-driven fluoropyrimidine resistance, linking SHIP2 expression to the adaptive upregulation of TYMS in breast cancer cells.

### SHIP2 rewires TYMS regulation from p53-dependent to β-catenin-dependent control

Our data above demonstrate that overexpression of either wild-type (WT) or phosphatase-dead (PD) SHIP2 enhances both basal and stress-induced accumulation of TYMS protein. In addition, we have shown that SHIP2 depletion modulates p53 transcriptional activity in a gene- and cell-type-specific manner (**Supplementary Figure 4**). We therefore examined the potential role of p53 in the context of SHIP2 overexpression. To address this question, we generated stable MDA-MB-231 cell lines expressing either a control shRNA or a p53-targeting shRNA in parental cells or in cells stably overexpressing WT or PD SHIP2. As shown in Figure 6 A, stable depletion of p53 in parental cells led to only a moderate decrease in active β-catenin levels. However, TYMS protein levels were significantly reduced under both basal and FuDR-induced stress conditions in these cells. These findings are consistent with our previous results, which showed that p53 depletion alters TYMS protein levels in breast cancer cells [16]. In contrast, p53 depletion in cells overexpressing WT or PD SHIP2 leads to a moderate increase in active β-catenin levels. Surprisingly, in SHIP2-overexpressing cells, p53 depletion resulted in only a modest reduction in basal and FuDR-induced TYMS protein levels compared with parental cells (**Figure 6 A-C**). These results suggest that stable SHIP2 overexpression promotes TYMS expression, at least in part, by activating β-catenin signaling, thereby reducing the dependence of TYMS regulation on p53. To further test this hypothesis, WT and phosphatase-dead SHIP2-overexpressing cells (all expressing shRNA against p53) were treated with a β-catenin inhibitor. As shown in Figure 6 D-E, inhibition of β-catenin markedly reduced TYMS protein levels in SHIP2-overexpressing cells, confirming that SHIP2-driven TYMS expression relies predominantly on β-catenin activity when p53 is depleted. Finally, we investigated how these molecular changes affect long-term cell growth using colony formation assays. Combined WT or PD-SHIP2 overexpression with P53 depletion conferred resistance to long-term treatment with FuDR relative to P53 depletion alone. However, combined treatment of β-catenin inhibition and FuDR reversed the resistance conferred by SHIP2 overexpression, with a stronger effect in PD-SHIP2 cells. Indeed, P53-depleted cells overexpressing WT or PD SHIP2, which normally exhibit greater colony-forming ability under FuDR treatment, displayed reduced survival and growth when β-catenin was concurrently inhibited (**Figure 6 F-G**). Collectively, these data indicate that SHIP2 overexpression functionally rewires TYMS regulation, shifting it from a p53-dependent mechanism in parental cells to a β-catenin–driven program in SHIP2-overexpressing cells.

**Figure 6:**
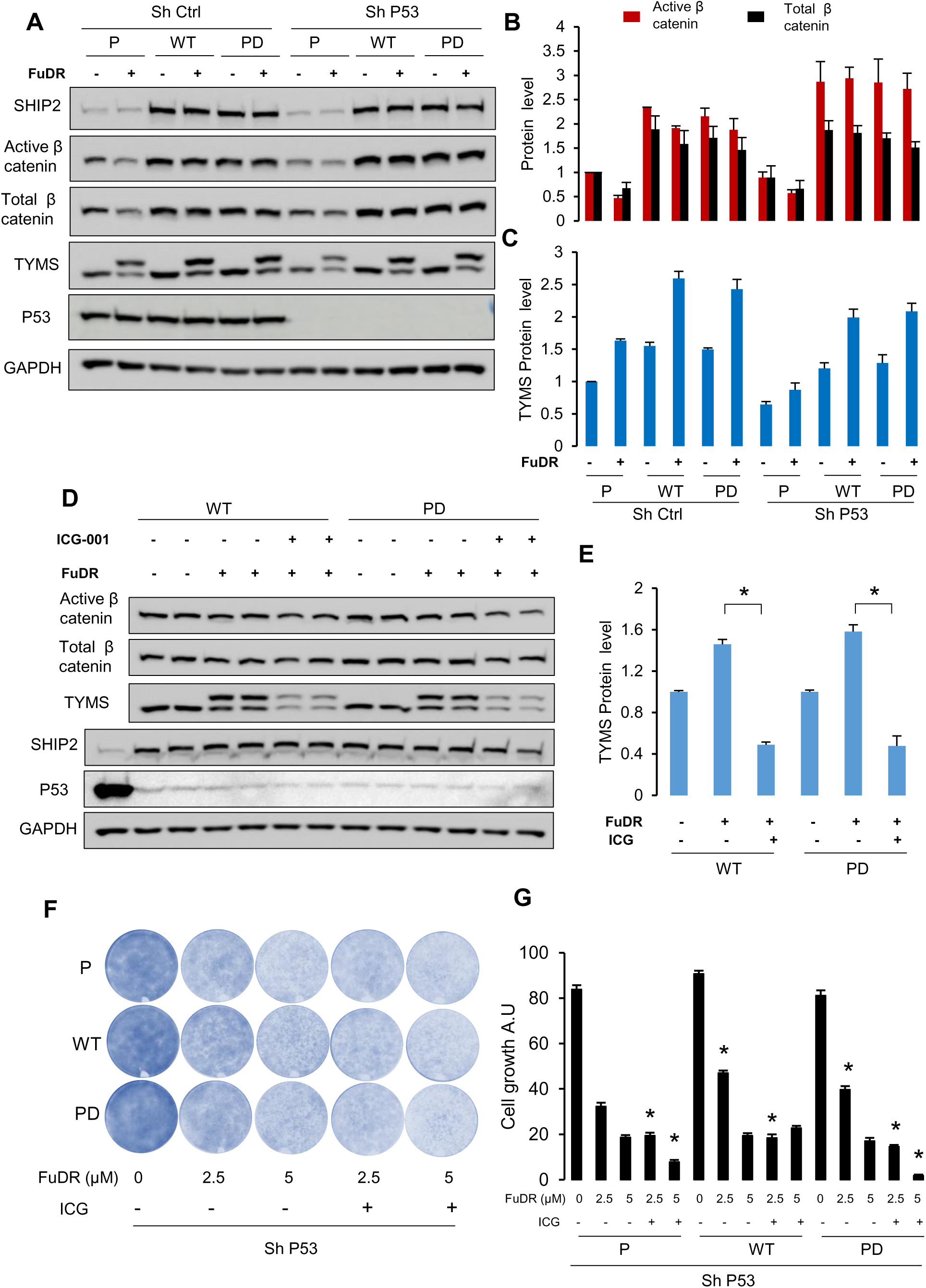
**(A)** Lysates from MDA-MB-231 parental cells or stably overexpressing WT or PD SHIP2, shRNA control, or stably expressing and shRNA against P53 treated or not with 50 µM FuDR for 24 hours, then stained with the indicated antibodies. n=3 biological replicates per condition. (**B-C**) Bar graphs showing the results of densitometric analysis of active, total β-catenin, and TYMS levels under the conditions obtained in A. The data are presented as the means ±SEMs. ANOVA, multiple comparisons: TYMS: F (11, 24) = 41.6, 6.4E-13. (**D**) Lysates from MDA-MB-231 cells stably overexpressing WT or PD SHIP2, stably expressing ShRNA against P53, treated or not with 50 µM FuDR for 24 hours, supplemented or not with 10µM β-catenin inhibitor (ICG-001), then stained with the indicated antibodies. n=4 biological replicates per condition. (**E**) Bar graphs showing the results of densitometric analysis of TYMS levels under the conditions obtained in D. The data are presented as the means ±SEMs. ANOVA, multiple comparisons: Tukey’s multiple comparisons test. Dunnett test. WT cells: F (2, 9) = 240.9, P<0.0001. PD cells: F (2, 9) = 65.8, P<0.0001. (**F-G**) Clongenic assay of MDA-MB-231 parental, WT, and PD SHIP2 cells stably expressing ShRNA against P53 treated with the indicated concentrations of FuDR, supplemented or not with 1 µM β-catenin inhibitor ICG-001 (ICG). Representative images from each condition were shown. ANOVA, multiple comparisons: F (20, 105) = 226.5, P=6.6E-77.

### SRC activity is required for SHIP2-dependent TYMS expression and nuclear β-catenin translocation

Our data indicate that the β-catenin-TYMS signaling axis regulated by SHIP2 is independent of its lipid phosphatase activity, suggesting that SHIP2 may function through protein-protein interactions. Previous studies have shown that SHIP2 can act as an adaptor protein, recruiting Src family kinases to FGFR receptor complexes, thereby sustaining ERK signaling [6]. Given the established role of SRC in regulating β-catenin stability and nuclear activity [26–27], we hypothesized that SRC protein could be the mediator via which SHIP2 modulates the β-catenin-TYMS axis.

To test this hypothesis, we first examined the impact of SHIP2 expression on SRC protein levels. As shown in Figure 7A, SHIP2 overexpression in MDA-MB-231 cells resulted in a marked increase in SRC protein levels, whereas SHIP2 depletion in both MCF7 and MDA-MB-231 cells led to reduced SRC expression (**Figure 7 A-B**). Consistently, SHIP2 overexpression enhanced nuclear accumulation of SRC, whereas SHIP2 knockdown decreased nuclear SRC levels (**Figure 4 G-H**, **Figure 5 A-B**). Consistent with our previous findings in liver cancer cells [28], co-immunoprecipitation analyses further demonstrated a physical interaction between SHIP2 and SRC, supporting a conserved functional association between these two proteins (**Figure 7C**). We next assessed whether SRC was required for TYMS regulation downstream of SHIP2. Transient knockdown of SRC using ShRNA significantly reduced TYMS protein levels under basal conditions in both control and SHIP2-overexpressing cells (**Figure 7 D**). Importantly, SRC downregulation also abrogated the FuDR-induced increase in TYMS expression, indicating that SRC activity is required for adaptive TYMS upregulation under chemotherapeutic stress (**Figure 7 E**). Collectively, these findings support a model in which SHIP2 promotes TYMS expression and β-catenin activity through a non-catalytic adaptor function involving SRC kinase.

**Figure 7:**
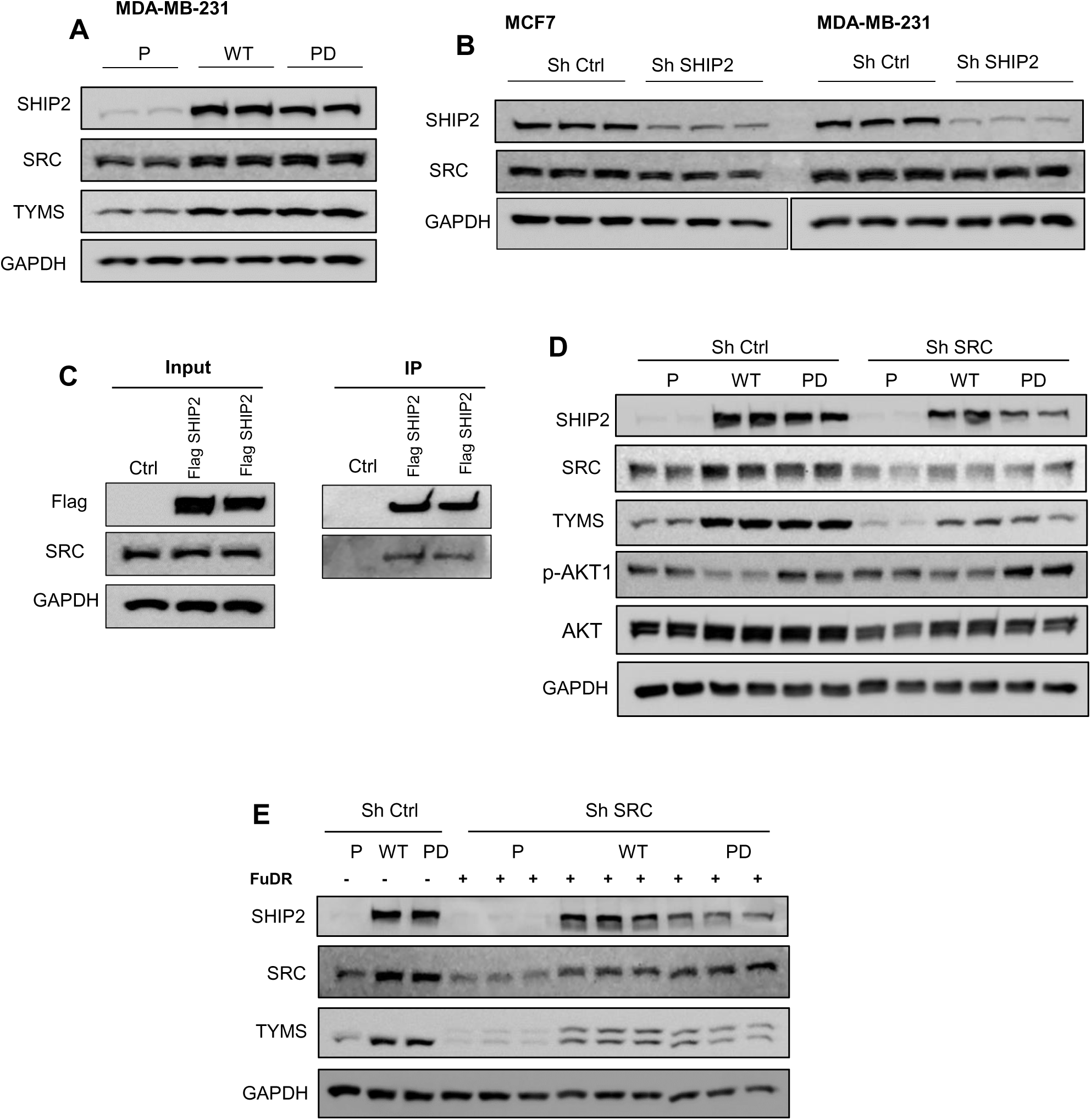
**(A-B)** Western blot analysis of c-SRC protein levels in MDA-MB-231 parental cells and cells overexpressing WT or PD SHIP2 (A) or in MDA-MB-231 and MCF7 cells stably expressing shRNA control or shRNA against SHIP2 (B). Representative of 3-5 biological replicates per condition. (**C**) Immunoprecipitation (IP) results showing the interaction between SHIP2 and c-SRC. Lysates from HEK293 transfected with Flag-SHIP followed by immunoprecipitation with anti-FLAG. Input and IP sample analyzed by immunoblot for the indicated antibodies. (**D-E**) Lysates from MDA-MB-231 parental cells or stably overexpressing WT or PD SHIP2, transiently transfected with shRNA control or ShRNA against SRC, treated or not with 50 µM FuDR for 24 hours, then stained with the indicated antibodies. Representative of 3 biological replicates per condition.

### Clinical association of SHIP2 expression with TYMS levels and patient outcome

Using the TCGA cancer database [15], our data above show that SHIP2 RNA expression levels positively correlate with TYMS mRNA levels, supporting the relevance of SHIP2-dependent TYMS regulation in our experimental model. To determine whether SHIP2 expression is altered in human breast cancer, we next analyzed an independent publicly available breast cancer proteomic dataset [29]. As shown in Figure 8 A, SHIP2 protein expression was significantly elevated across all major breast cancer subtypes compared with normal breast tissue. Interestingly, SHIP2 expression was further increased in tumors exhibiting alterations in WNT signaling components, consistent with our in vitro findings linking SHIP2 expression to changes in β-catenin signaling activity (**Figure 8 B**). To assess the clinical significance of SHIP2 expression, we performed Kaplan–Meier survival analyses [30]. Elevated INPPL1 expression alone was not significantly associated with reduced overall survival (HR = 1.09, 95% CI 0.90–1.31, p = 0.40) (**Figure 8 C**). However, patients with high SHIP2 expression exhibited significantly poorer relapse-free survival and distant metastasis-free survival compared with patients expressing low levels of SHIP2 (**Figure 8 D-E**). These findings suggest that while SHIP2 expression may not substantially affect overall mortality, it is associated with disease recurrence and metastatic progression.

**Figure 8:**
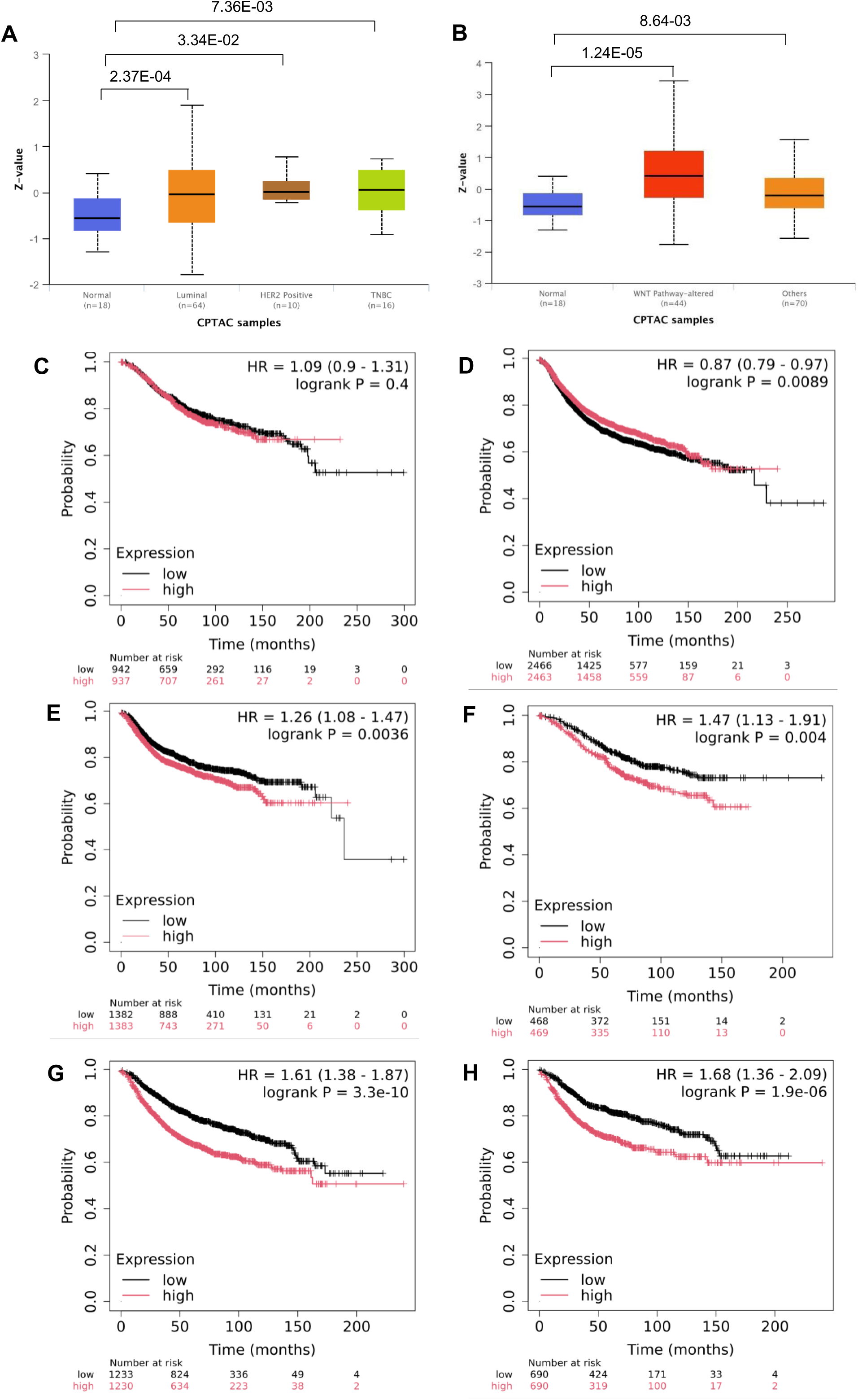
(**A**) Plot showing SHIP2 protein expression between normal breast tissue and breast tumor tissue subtype. (**B**) Plot showing SHIP2 protein expression between normal and breast cancer based on WNT pathway status. (**C-E**) Kaplan-Meier curves for survival in patients with breast cancer in relation to SHIP2 expression: (C) overall survival, (D) Relapse-Free Survival, and (E) Distant Metastasis-Free Survival. (**F-H**) Kaplan-Meier curves for survival in a subset of patients with breast cancer with high SHIP2 and TYMS SHIP2 expression: (F) overall survival, (G) Relapse-Free Survival, and (H) Distant Metastasis-Free Survival.

Given the strong association between SHIP2 and TYMS expression, we next examined the prognostic impact of their combined expression. Strikingly, patients whose tumors showed high expression of both INPPL1 and TYMS had significantly worse clinical outcomes than those with low expression of one or both genes. Indeed, combined high SHIP2/TYMS expression was associated with reduced overall survival, relapse-free survival, and distant metastasis-free survival, identifying a subgroup of patients with particularly aggressive disease (**Figure 8 F-H**). These findings support a clinically relevant SHIP2-TYMS signaling axis and suggest that coexpression of these genes may serve as a prognostic biomarker for poor outcome in breast cancer.

Collectively, these clinical data corroborate our mechanistic studies and indicate that elevated SHIP2 expression is associated with increased TYMS expression, dysregulated WNT/β-catenin signaling, and adverse patient outcomes. The particularly poor prognosis observed in tumors co-expressing high levels of SHIP2 and TYMS further supports the biological and clinical significance of the SHIP2–TYMS regulatory pathway in breast cancer progression.

## DISCUSSION

Fluoropyrimidine resistance remains a major limitation in the treatment of solid tumors, and adaptive upregulation of TYMS is a central mechanism underlying this process. The upstream signaling pathways that control TYMS expression in response to chemotherapeutic stress remain incompletely understood. In this study, we identify the lipid phosphatase SHIP2 as a previously unrecognized regulator of TYMS expression and fluoropyrimidine response in breast cancer cells. Our findings reveal that SHIP2 promotes TYMS transcription and protein accumulation independently of its lipid phosphatase mechanism, which involves SRC and β-catenin signaling. Moreover, we demonstrate that SHIP2 functionally rewires TYMS regulation from a TP53-dependent stress response to a β-catenin-driven program, thereby sustaining TYMS expression and promoting resistance to Floxuridine.

A key finding of our study is that SHIP2 regulates TYMS independently of its lipid phosphatase activity. Although SHIP2 is classically defined as a negative regulator of PI3K/AKT signaling, both wild-type and phosphatase-dead SHIP2 similarly increased TYMS expression, indicating that this function does not rely on its catalytic activity. Our findings are consistent with a previous study describing the adaptor function of SHIP2 in growth factor signaling, which shows that it facilitates the assembly of signaling receptor complexes [6]. In this context, our data support a model in which SHIP2 acts as an adaptor protein that promotes oncogenic signaling independently of PI3K/AKT modulation.

Mechanistically, we identify SRC as a critical intermediate protein linking SHIP2 to β-catenin activation. Indeed, SHIP2 overexpression increased SRC protein levels and nuclear localization, whereas SRC downregulation reduced TYMS expression. These findings are consistent with established roles of SRC in regulating β-catenin activity and localization, which can promote its dissociation from adherens junctions and enhance its nuclear signaling [26]. Our results, therefore, position SRC as a key mediator of SHIP2-driven β-catenin activation.

We further show that β-catenin is required for SHIP2-mediated TYMS expression and chemoresistance. Indeed, pharmacological inhibition of β-catenin transcriptional activity suppressed TYMS induction and reversed resistance to Floxuridine in SHIP2-overexpressing cells. Interestingly, inhibition of β-catenin/CBP interaction, but not β-catenin/TCF interaction, was sufficient to block TYMS upregulation, suggesting that TYMS is regulated through a non-canonical β-catenin transcriptional program. These findings expand the repertoire of β-catenin-dependent transcriptional outputs and suggest that cofactor selection may determine context-specific gene regulation. In line with this hypothesis, independent studies indicate that under conditions of cellular stress, β-catenin can engage alternative transcriptional partners, including FOXO family transcription factors and chromatin-modifying enzymes, thereby regulating gene expression programs distinct from canonical Wnt targets. These stress-adaptive β-catenin complexes have been implicated in cell survival, redox regulation, and therapy resistance [31–33].

Another important finding in our study is the identification of a shift in the regulatory mechanisms controlling TYMS expression. Indeed, our data indicate that TYMS is subject to dual regulation by P53 and β-catenin pathways. In MDA-MB-231 parental cells, TYMS expression is at least partly dependent on p53, as its depletion significantly reduces TYMS levels. In contrast, in SHIP2 overexpressing cells, TYMS expression becomes less sensitive to p53 loss and shows increased dependence on β-catenin activity. These findings suggest that SHIP2 promotes a redistribution of regulatory control from a predominantly p53-associated mechanism toward a β-catenin-driven program, rather than a complete switch. Such a shift may allow tumor cells to maintain TYMS expression and sustain nucleotide synthesis under conditions of genotoxic stress or impaired p53 function.

Although this was not investigated in detail in the present study, we observed a similar pattern in SRC expression. Indeed, p53 depletion reduced SRC protein levels in parental cells; however, this effect was markedly attenuated in SHIP2-overexpressing cells (**Supplementary Figure 6**). These observations suggest that high SHIP2 expression may extend this regulatory shift upstream, partially uncoupling SRC expression from p53-dependent regulation. Given the role of SRC in promoting β-catenin stability and activity, this mechanism may further reinforce β-catenin-dependent regulation of TYMS in SHIP2-overexpressing cells.

It is worth noting that our findings are primarily based on experiments performed in MDA-MB-231 cells that express a mutant p53. Due to the inability to generate stable SHIP2-overexpressing MCF7 cells, we were unable to directly assess whether a similar regulatory switch occurs in a wild-type p53 context. This limitation suggests that the extent and nature of TYMS rewiring may depend on cellular context, particularly p53 status. Nevertheless, our data support a model in which SHIP2 promotes a shift toward β-catenin-dependent transcriptional control of TYMS, potentially enabling tumor cells to sustain nucleotide synthesis and proliferative capacity under conditions of genotoxic stress or impaired p53 function.

Our findings also show that the effect of SHIP2 depletion on fluoropyrimidine sensitivity is context dependent. While SHIP2 knockdown sensitized MCF7 cells to Floxuridine, this effect was less pronounced in MDA-MB-231 cells. This difference may be linked to higher basal TYMS levels in MDA-MB-231 cells, which remain above a functional threshold required to sustain nucleotide synthesis even after SHIP2 depletion. These observations suggest a threshold model in which the impact of SHIP2 targeting depends on baseline TYMS abundance and the metabolic state of tumor cells. In this context, cells with high TYMS levels may retain sufficient enzymatic activity to buffer the effects of SHIP2 loss, thereby attenuating the impact of changes in SHIP2 levels on sensitization to fluoropyrimidine treatment. Further studies will be required to understand this threshold model and determine its relevance across different tumor types and therapeutic contexts. Analysis of clinical datasets supports the translational relevance of SHIP2-TYMS signaling axis. Indeed, SHIP2 expression positively correlates with TYMS levels in breast cancer cohorts and is associated with poorer patient outcomes, particularly in terms of disease recurrence and metastatic progression. These observations are consistent with our experimental findings and suggest that elevated SHIP2 expression may contribute to therapeutic resistance in human tumors.

Despite these findings, our study has some limitations. First, our study is primarily based on in vitro models, and in vivo validation will be required to fully establish the role of SHIP2 in drug resistance and metastasis. Second, although our data support a transcriptional role for β-catenin in regulating TYMS, the precise promoter elements and transcriptional complexes involved remain to be defined. Therefore, future studies using chromatin-based approaches will be necessary to elucidate the direct regulatory mechanisms. Finally, the potential contribution of additional signaling pathways affected by changes in SHIP2 levels cannot be excluded.

In conclusion, our study identifies SHIP2 as a non-canonical regulator of TYMS expression and fluoropyrimidine resistance. By promoting SRC-dependent activation of β-catenin, SHIP2 sustains TYMS expression and reprograms its regulation away from p53-dependent stress signaling. These findings uncover a signaling mechanism that links oncogenic signaling to nucleotide metabolism and suggest that targeting the SHIP2–SRC–β-catenin axis may represent a therapeutic strategy to overcome fluoropyrimidine resistance.

## Materials and Methods

### Reagents and chemical inhibitors

Dulbecco’s Modified Eagle Medium (DMEM, high glucose, 4.5 g/L glucose, supplemented with L-glutamine and sodium pyruvate) was obtained from Capricorn Scientific (Cat# DMEM-HPA). Fetal bovine serum (FBS) was purchased from Sigma-Aldrich, and penicillin-streptomycin solution was obtained from Thermo Fisher Scientific (Cat# 15140122). The following pharmacological agents were used: 5-fluorouracil (Cat# F6627-1G), 5-fluoro-2′-deoxyuridine (Cat# L16497. ME), PNU-74654 (Medchemexpress, #HY-101130), ICG-001 (Medchemexpress, #HY-14428). Stock solutions were prepared according to the manufacturers’ recommendations and diluted in culture medium immediately prior to use. Vehicle-treated controls received the corresponding DMSO concentration.

### Cell culture and treatment conditions

HEK293T, MCF7, and MDA-MB-231 cells were provided by Dr. Anna Marusiak (IMol Institute). Cells were maintained in DMEM supplemented with 10% FBS, 100 U/mL penicillin, and 100 U/mL streptomycin. Cultures were incubated at 37 °C in a humidified atmosphere containing 5% CO₂ and were routinely passaged to maintain exponential growth. Drug treatments were performed at the concentrations and durations indicated in the figure legends.

### Protein extraction and Western blot analysis

Following each treatment, cells were washed once with cold PBS and lysed in buffer containing 50 mM Tris-HCl (pH 8.0), 150 mM NaCl, 1.0% NP-40, 0.1% SDS, benzonase (25 U/mL), PMSF (100 µM), and 5% glycerol. Cell lysates were incubated on ice and subsequently clarified by centrifugation at 12,000 × g for 10 min at 4 °C. Protein concentrations were measured using a BCA protein assay. Equal amounts of total protein (10-20 µg) were separated by SDS-PAGE and transferred onto nitrocellulose membranes (Cat# 10600004). Upon transfer, membranes were stained with ponceau red to verify transfer quality, then blocked for 1 h in TBST containing 5% non-fat milk and incubated overnight at 4 °C with primary antibodies. After washing, membranes were incubated with HRP-conjugated secondary antibodies for 1 h at room temperature. Immunoreactive bands were visualized using the Amersham ImageQuant 800 detection system. Densitometric analysis was performed using ImageJ software. For measuring total TYMS protein level, both slower- and faster-migrating bands were included in the calculation.

### Lentiviral shRNA production

Lentiviral particles were produced as previously described [34]. HEK293T cells were first seeded in 6-well plates at approximately 60% confluency. Prior to transfection, the growth medium was replaced with fresh and complete DMEM. Packaging (psPAX2), envelope (VSV-G), and shRNA expression plasmids were diluted in Opti-MEM and combined with PEI diluted separately in Opti-MEM. After incubation at room temperature to allow complex formation, the transfection mixture was added dropwise to the cells. The medium was replaced after 4 hours. Viral supernatants were collected 48 hours post-transfection, cleared by centrifugation at 12,000 × g for 5 min, and stored at -80 °C until use. The following shRNAs targeting SHIP2 were used: SH1 (TRCN0000052808), SH2 (TRCN0000052809). ShRNA control Empty (Sigma Aldrich, # SHC001), P53 (shp53 pLKO.1 puro: Addgene, #19119).

### Lentiviral infection and antibiotic selection

MCF7 and MDA-MB-231 cells were seeded in 6-well plates and infected with lentiviral supernatants mixed 1:1 with fresh culture medium. Cells were incubated with the virus overnight, after which the medium was replaced. Forty-four hours post-infection, puromycin selection was carried out (3 µg/mL for MCF7 and 2 µg/mL for MDA-MB-231) and continued for 96 hours. Stable knockdown was confirmed by Western blotting and quantitative real-time PCR.

### RNA isolation

Total RNA was isolated using TRI Reagent according to the manufacturer’s instructions. Briefly, cells were lysed in TRI Reagent and incubated at room temperature for 5 minutes. The mixture was phase-separated with chloroform. Samples were centrifuged at 15,000 × g for 15 min at 4 °C, and the aqueous phase containing RNA was transferred to a new tube. RNA was precipitated with isopropanol, washed with 75% ethanol, air-dried, and resuspended in RNase-free water. RNA concentration and purity were assessed prior to cDNA synthesis.

### cDNA synthesis and quantitative real-time PCR

cDNA synthesis was performed using 1 µg of total RNA and a high-capacity cDNA reverse transcription (Thermo Fisher Scientific, Cat#4368814), following manufacturer instructions. Quantitative PCR was performed using PowerUp™ SYBR™ Green Master Mix (Applied Biosystems, Cat# A25776) on a LightCycler 480 real-time PCR system. Each reaction contained 10 ng cDNA and 300 nM forward and reverse primers and was run in technical duplicates. Cycling conditions consisted of an initial activation step at 95 °C, followed by 40 amplification cycles. Melting curve analysis was performed to verify amplification specificity. Gene expression levels were calculated using the ΔΔCt method and normalized to Gapdh. Primer sequences are listed below.

Forward-TYMS-CTGCTGACAACCAAACGTGTG

Reverse-TYMS-GCATCCCAGATTTTCACTCCCTT

Forward-GAPDH-GGAGCGAGATCCCTCCAAAAT

Reverse-GAPDH-GGCTGTTGTCATACTTCTCATGG

Forward-CDKN1A- CGATGGAACTTCGACTTTGTCA

Reverse-CDKN1A- GCACAAGGGTACAAGACAGTG

Forward-GADD45A- CCCTGATCCAGGCGTTTTG

Reverse-GADD45A- GATCCATGTAGCGACTTTCCC

Forward- SESN2-CCTCTGGGCGAGTAGACAAC

Reverse- SESN2-GGAGCCTACCAGGTAAGAACA

Forward-TP53-GAGGTTGGCTCTGACTGTACC

Reverse-TP53-TCCGTCCCAGTAGATTACCAC

Forward-BIRC5-AGGACCACCGCATCTCTACAT

Reverse-BIRC5-AAGTCTGGCTCGTTCTCAGTG

Forward-Inppl1-ATGACCGGGATGCCTCAGAT

Reverse-Inppl1-CCTTTCAGGTAGTCGTGCGAG

### Cell cycle analysis

To examine the impact of 5-fluoro-2′-deoxyuridine on the cell cycle, MDA-MB-231 cells were treated for 72 h, harvested by trypsinization, and washed with PBS. Cells were then fixed in 70% ethanol at −20 °C for at least 24 hours. Propidium iodide staining was performed as follows: Fixed cells were collected by centrifugation at 20,000 × g for 25 min at 4 °C. Cell pellets were re-suspended in PBS, then collected by centrifugation at 20,000 × g for 25 min at 4 °C. Cells were incubated in PBS supplemented with RNase A and propidium iodide (Invitrogen, #P1304MP) (50µg/ml) for 30 min at room temperature. DNA content was then analyzed using a NovoCyte flow cytometer, and cell cycle distribution was quantified using the NovoExpress software package.

### Colony formation assay

MCF7 and MDA-MB-231 cells stably overexpressing WT SHIP2, PD SHIP2, or a shRNA targeting SHIP2, or a shRNA control, were seeded into 24-well plates (2500 cells/well). After 24 hours, cells were treated with vehicle or the indicated concentrations of each inhibitor and cultured for 12-16 days under standard conditions. At the end of each experiment, colonies were first fixed with 100% methanol and then stained with crystal violet. After drying, stained colonies were imaged using GelDoc (Bio-Rad). Cell colonies were then solubilized in 10 % acetic acid, and the absorbance of the solution was measured at 590 nm using a plate reader (Multiskan™ FC Microplate Photometer, Thermo Scientific™).

### Nuclear and cytosolic fractionation

Control and treated cells (in a 10 cm culture dish) were first washed with cold PBS. Cells were then collected into 1.5 mL microcentrifuge tubes using a plastic cell scraper and cold PBS. Cells were pelleted by centrifugation at 2,500 rpm for 3 min at 4 °C. The cell pellets were resuspended in 1 mL of ice-cold PBS containing 0.1% NP-40 (Sigma-Aldrich). After five rounds of trituration using a P1000 micropipette, nuclear and cytosolic fractions were further separated by mechanical disruption with a Dounce homogenizer using a tight-fitting pestle (12 strokes). The homogenate was centrifuged at 10,000 × g for 3 min at 4 °C. The supernatant, containing the cytosolic fraction, was transferred to a fresh tube. The pellet, containing the nuclear fraction, was washed by resuspension in 1 mL of ice-cold PBS and centrifuged again at 10,000 × g for 3 min at 4 °C. The supernatant was discarded. The nuclear pellet was then resuspended in protein lysis buffer (described above) and sonicated using a probe sonicator. After protein quantification, equal amounts of protein from each fraction and condition were used for Western blot analysis.

### Co-immunoprecipitation

Co-immunoprecipitation was carried out as described previously [28]. Control and transfected HEK293 cells expressing Flag-SHIP2 were allowed to grow for 24 hours. Cells were washed with cold PBS and directly collected using immunoprecipitation buffer (IP-Buffer) containing 20 mM Tris-HCl (pH 7.4), 150 mM NaCl, 1 mM EGTA, 0.5% Igepal, 10% glycerol, and PMSF. Lysates were homogenized by agitation at 4 °C for 5 minutes (20 rpm/minute), then centrifuged (10000 g, 10 min at 4 °C). Protein concentration was determined using a BCA protein assay kit (Thermo Scientific, #23225). Equal amounts of cell lysate were then incubated with the anti-Flag antibody (Sigma Aldrich, #F1804) for 4 hours at 4 °C on a rotating platform. Protein A Agarose beads (Thermo Fisher Scientific, #20333) were used to collect the immuno-complex. After two washes with cold PBS, beads were added to the lysate-antibody and incubated for 1 h at 4 °C on a rotating platform. After three washes with PBS, beads were resuspended in 1XSDS gel loading buffer (50 mM Tris-HCl, pH 6.8, 2% SDS, 10% glycerol, 1% β-mercaptoethanol, 12.5 mM EDTA, 0.02% bromophenol blue) and stored at -80 °C until further use. CO-IP and input samples were analyzed by standard western blotting.

## Statistical analysis

All quantitative data are presented as mean ± SEM from at least three independent biological replicates. Statistical analyses were performed using GraphPad Prism software. Differences between the two groups were evaluated using unpaired two-tailed tests, while comparisons among multiple groups were performed using one-way ANOVA followed by Dunnett’s post hoc test. Statistical significance was defined as P < 0.05.

## DECLARATIONS

The AI tools Grammarly and ChatGPT were used only at the final stage of manuscript preparation, solely for English corrections and clarifications.

## Ethics approval and consent to participate

not applicable.

## Consent for publication

not applicable.

## Availability of data and materials

The data generated during the current study are available from the corresponding author on reasonable request.

## Authors’ contributions

Conceptualization, A.A., Methodology; A.A., Investigation; A.A. and A.E.S., original draft preparation, AA; Writing review and editing, A.A.; Funding acquisition, A.A.

---

**Supplementary Figure 1:**
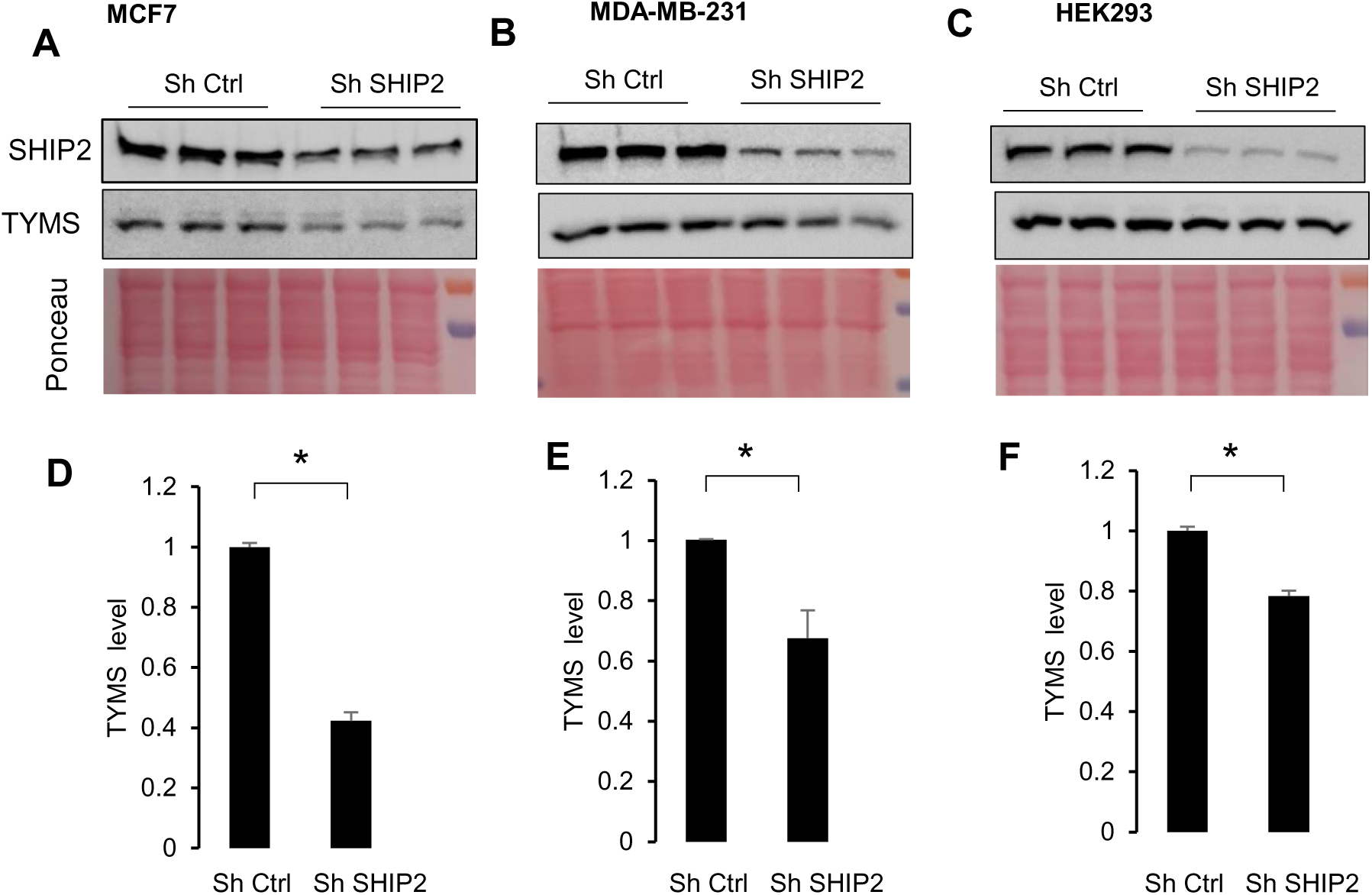
SHIP2 depletion alters TYMS RNA and protein levels. (**A-C**) Lysates of MCF7, MDA-MB-231, and HEK293 cells stably expressing Sh control or small hairpin against SHIP2 were stained with the indicated antibodies. (**D-F**) Bar graphs showing the results of densitometry analysis of TYMS protein levels under the conditions obtained in A and C. The data are presented as the means ±SEMs. ANOVA, unpaired T-Test *P<0.005.

**Supplementary Figure 2:**
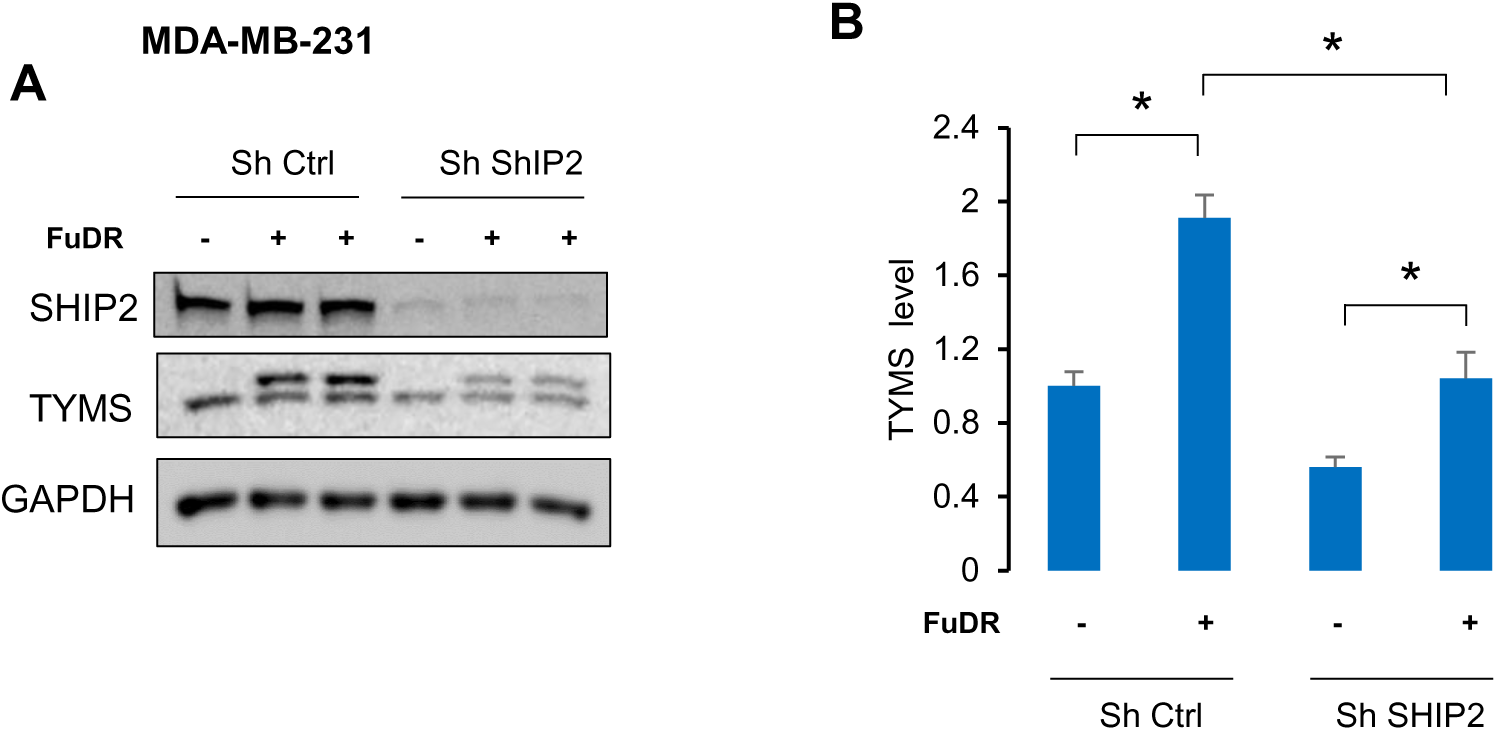
(**C**) Lysates of MDA-MB-231 cells stably expressing Sh control or a separate small hairpin (SH2) against SHIP2, treated or not with 50µM 5-FuDR for 24 hours, were stained with the indicated antibodies. (**D**) Bar graphs showing the results of densitometric analysis of TYMS protein levels obtained in A. The data are presented as the means ±SEMs. ANOVA, multiple comparisons: Dunnett test. F (3, 14) =19.2, P <0.001.

**Supplementary Figure 3:**
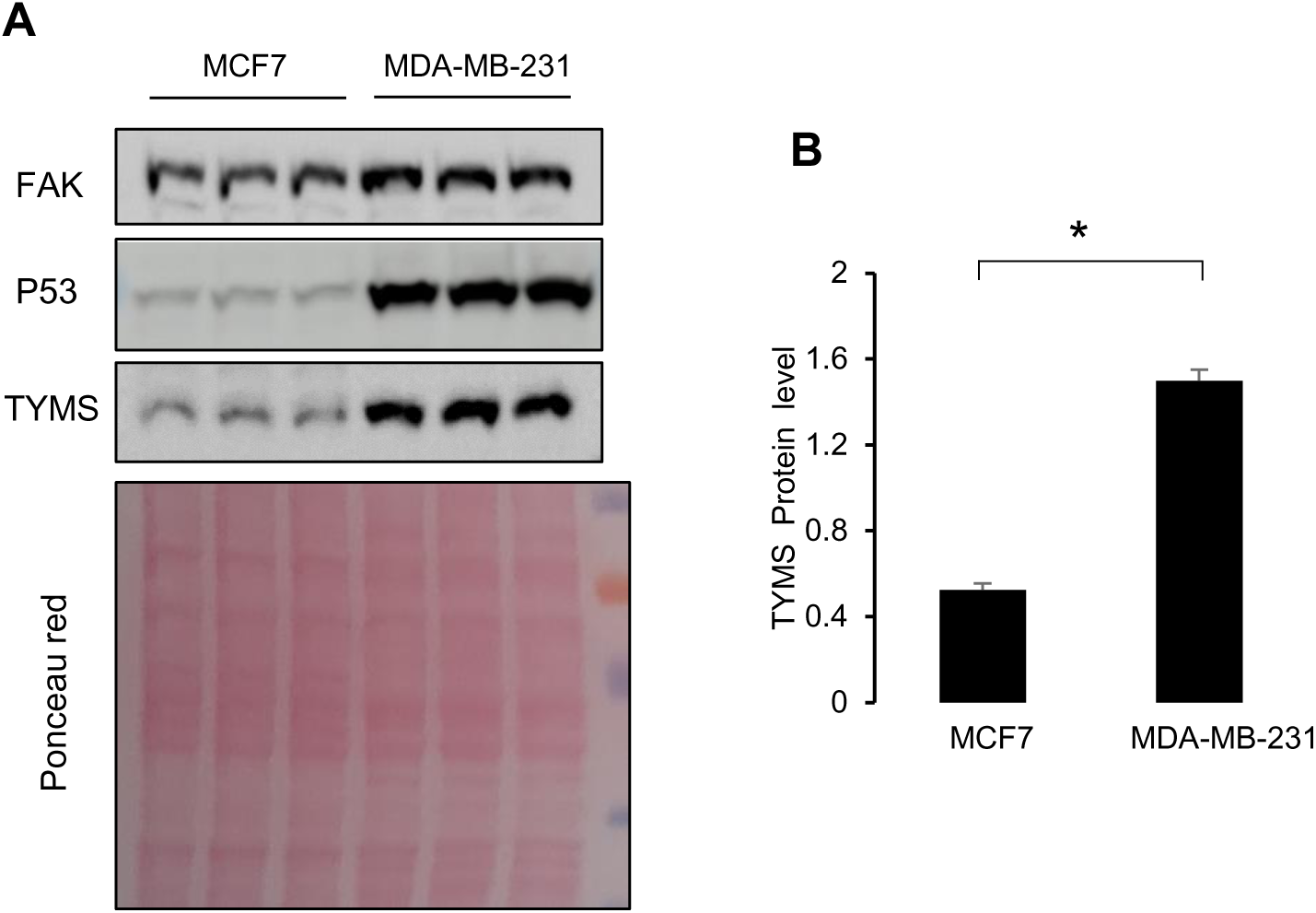
TYMS protein abundance is cell-type dependent. (**A**) Lysates of MCF7 and MDA-MB-231 WT cells stained with the indicated antibodies. n = 3 biological replicates per cell line. (**B**) Bar graph shown quantification of TYMS levels obtained in A. The data are presented as the means ±SEMs. Unpaired T-Test *P<0.005.

**Supplementary Figure 4:**
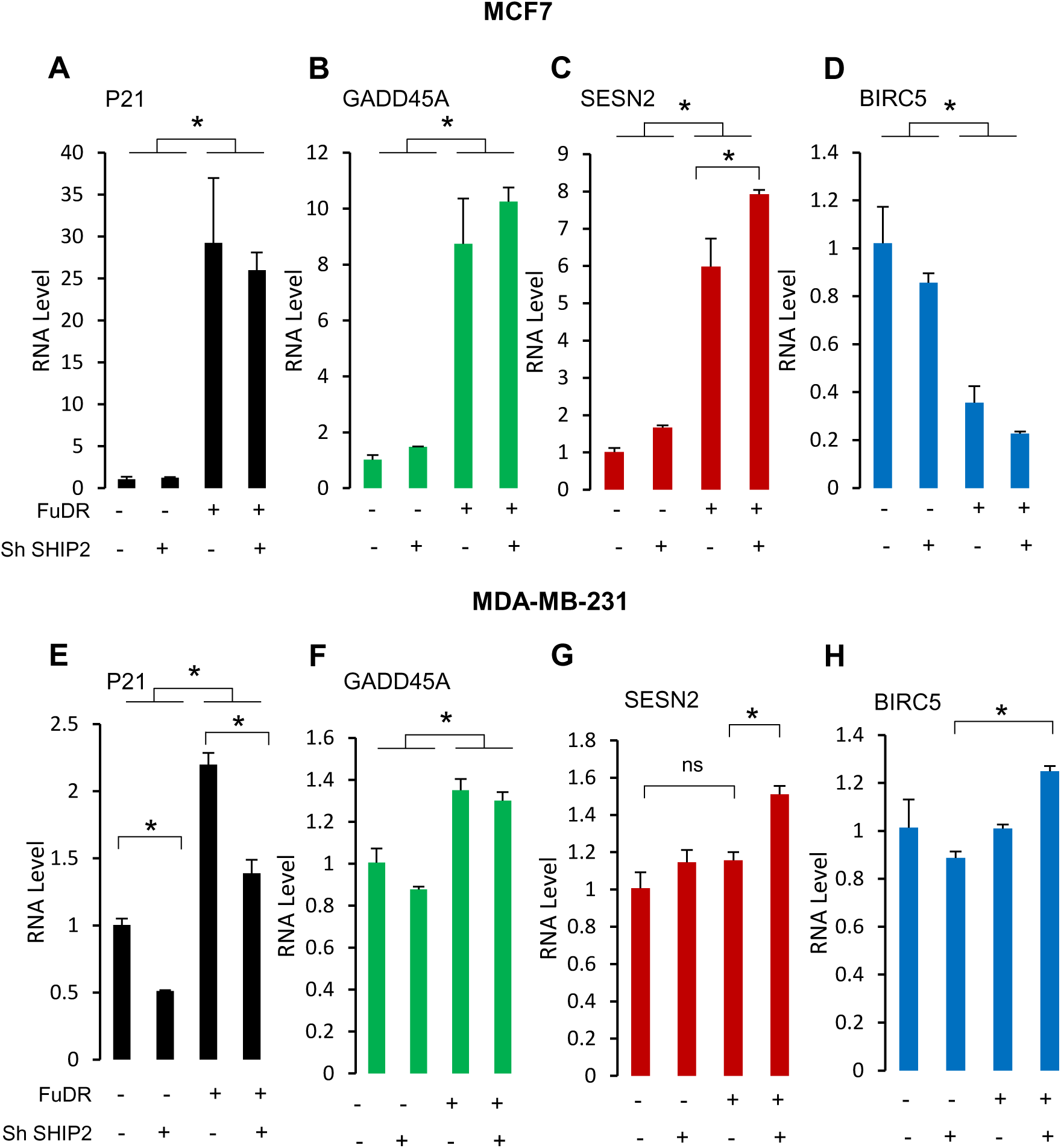
SHIP2 depletion alters stress-induced changes in P53 activity. (**A-H**) Bar graphs showing RNA levels of P53 targets and stress-inducible genes in MCF7 and MDA-MB-231 cells stably expressing Sh control or SH against SHIP2 treated or not with 50µM FuDR during 24 hours. The data are presented as the means ±SEMs. ANOVA, multiple comparisons: Dunnett test. **MCF7:** P21 F (3, 8) =14.58, p˂0.005. GADD45A F (3, 8) =31.9, p˂0.001. SESN2 F (3, 8) = 75.5, p ˂ 0.001. BIRC5 F (3, 8) =19.9, p˂0.001. **MDA-MB-231:** SESN2 F (3, 8) =11.8, p˂0.005. BIRC5 F (3, 8) =5.9, p˂0.01. P21 F (3, 8) =76.6, p˂0.0001. GADD45A F (3, 8) =16.96, p˂0.0001.

**Supplementary Figure 5:**
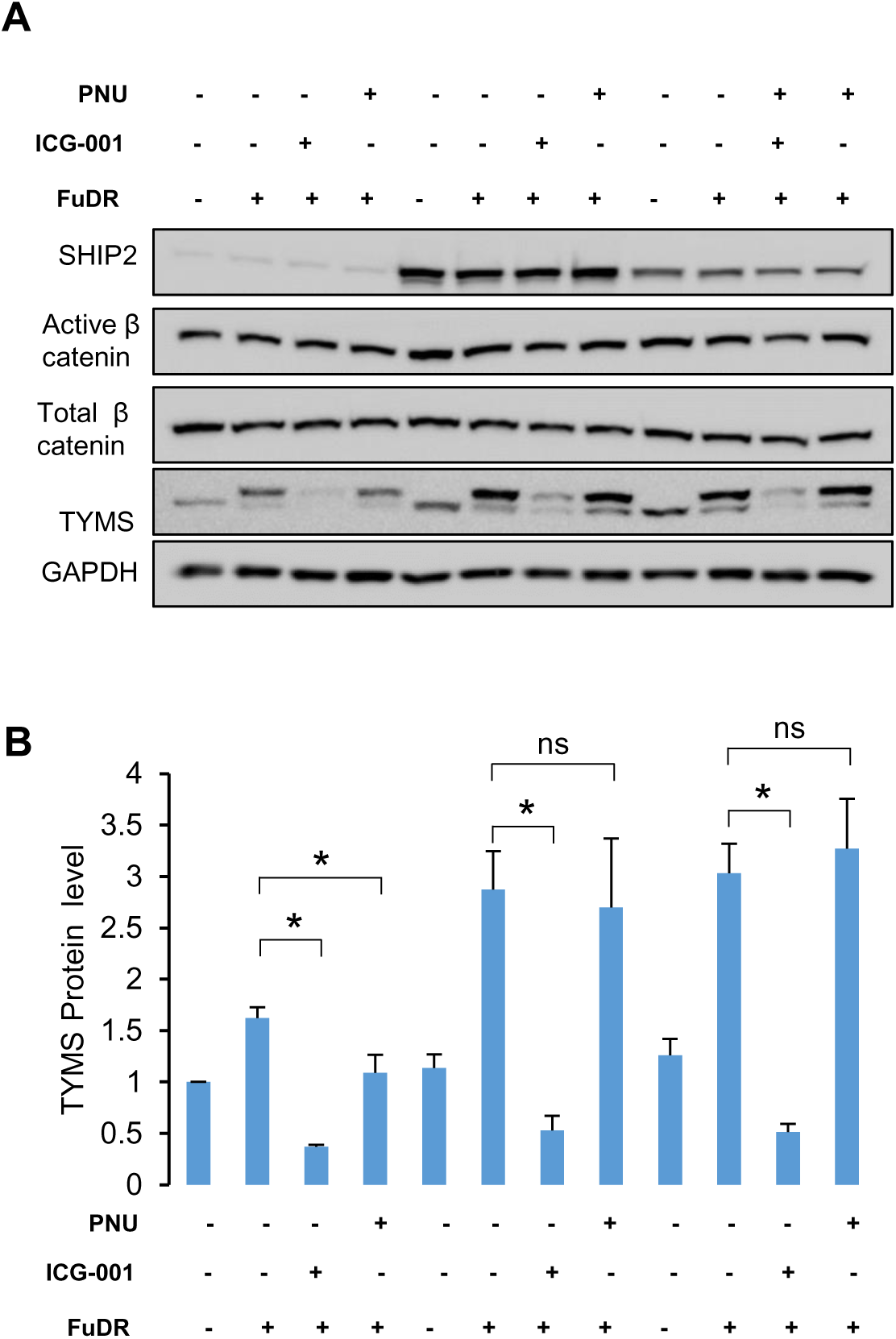
**(A)** Lysates from MDA-MB-231 parental cells or stably overexpressing WT or PD SHIP2 control or treated with 50 µM FuDR for 24 hours supplemented or not with 10 µM β-catenin inhibitors (ICG-001) or 20 µM PNU-74654, then stained with the indicated antibodies. n=3 biological replicates per condition. **(D)** Bar graphs showing the results of densitometric analysis of TYMS levels under the conditions obtained in A. The data are presented as the means ±SEMs. ANOVA, multiple comparisons: Dunnett test. Parental cells: F (3, 8) = 24.5, p˂0.005. WT Cells: F (3, 8) = 8.5, p˂0.005. PD Cells: F (3, 8) = 20.8, p˂0.005.

**Supplementary Figure 6:**
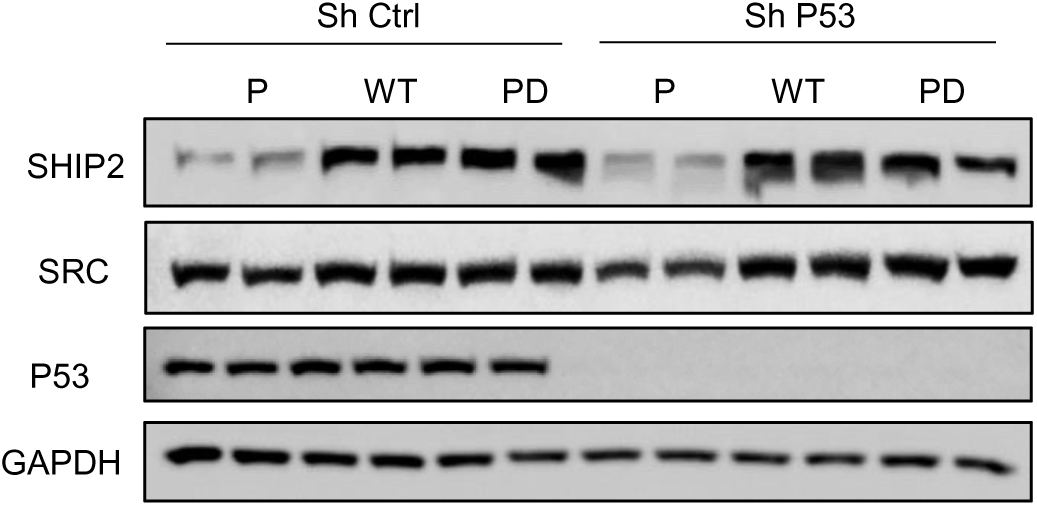
Lysate of MDA-MB-231 parental cells and cells overexpressing WT or PD SHIP2 stably expressing shRNA control or shRNA against P53 stained for the indicated antibodies.

